# Human Mendelian disease and *in vivo* mutagenesis screening define the molecular architecture of the U8 snoRNA

**DOI:** 10.1101/2025.10.10.681432

**Authors:** Andrew P. Badrock, Katryna Bell, Jian You Lau, Ankit Pathak, Cameron Wyatt, Christos Prapiadis, Raymond T. O’Keefe, Joseph A. Marsh, Grzegorz Kudla, Yanick J. Crow

## Abstract

Over 50 mutations in the vertebrate specific box C/D small nucleolar RNA U8 have been reported to cause leukoencephalopathy with calcifications and cysts (LCC), a progressive cerebral microangiopathy presenting at any age between early infancy and late adulthood. Notably, the majority of these disease-associated mutations are annotated as variants of uncertain significance due to an inability to predict their effect on U8 function. Here, using a zebrafish bioassay, we performed *in vivo* mutagenesis screening to explore the molecular pathology of LCC. We demonstrate that LCC-associated mutations cluster in function and molecular behaviour according to distinct structural domains within U8, differentially impairing U8 processing, activity and stability. Further, we show that when U8 function is hypomorphic, exogenous human U8 and the endogenous zebrafish U8.3 paralogue increase in stability through a newly discovered post-transcriptional mechanism. This latter response is compromised in patients with LCC due to the presence of a common polymorphism of an N^6^-methyladenosine modified nucleotide, which we predict to impact disease penetrance.

## Introduction

U8 is a vertebrate specific box C/D small nucleolar RNA (snoRNA) encoded by *SNORD118*, with biallelic mutations in *SNORD118* / U8 causing the Mendelian autosomal recessive disorder leukoencephalopathy with calcifications and cysts (LCC), also referred to as Labrune syndrome^1,2^. LCC is a specifically neurological condition affecting cerebral small blood vessels, and is characterised by the radiological triad of cerebral white matter disease, intracranial calcifications and cysts. LCC can present at any age from early infancy to late adulthood, and is associated with significant morbidity and premature mortality. Of particular note, although LCC is rare and inherited as an autosomal recessive trait, there is no enrichment for consanguinity and molecular homozygosity in patient cohorts. Indeed, in a recent comprehensive review, 93% of affected individuals were compound heterozygotes, likely carrying one ‘severe’ (null) and one ‘mild’ (hypomorphic) mutation^2^. This observation is extremely important, suggesting both that complete absence of U8 function due to homozygosity for two null mutations is incompatible with life, and that homozygosity for two hypomorphic mutations does not usually result in LCC. Given that *SNORD118* is non-protein encoding, a significant number of the LCC-associated mutations are annotated in the Genome Aggregation Database (gnomAD) v.4.1.0 as variants of uncertain significance.

Unlike most box C/D snoRNAs that mediate 2^′^-O-methylation of ribosomal RNA (rRNA), U8 functions to process the polycistronic precursor rRNA (pre-rRNA), facilitating the removal of internal transcribed spacer 2 and 3^′^-external transcribed spacer, essential steps in the biogenesis of 5.8S and 28S, components of the large ribosomal subunit^3^. Whereas mice and humans harbour a single copy of *Snord118 / SNORD118* (located on chromosomes 11 and 17, respectively), zebrafish have five copies of U8 situated on chromosome 10, presumably related to ancestral genomic duplication events. Zebrafish U8.3 is specifically expressed during early embryogenesis, before the other four copies of U8 (U8.1, U8.2, U8.4 and U8.5) are transcribed from approximately 2 days post fertilisation^4^. The role of U8 in 5.8S and 28S rRNA biogenesis was first defined in the *xenopus* oocyte^3^. Defects in ribosome biogenesis induce the tumour suppressor p53 (tp53)^5–7^, and we and others have shown that loss of U8 function results in activation of tp53 in mammalian cell lines^8^, zebrafish^4^ and cerebral organoids derived from human pluripotent stem cells expressing an LCC-causative U8 mutation^8,9^. Taken together, these models of U8 dysfunction indicate that LCC should be classified as a ribosomopathy, a heterogeneous class of human syndromes caused by mutations in factors required for the generation of functional ribosomes.

RNA polymerase II transcribes U8 as a 161-nucleotide precursor with a 3^′^ extension, with latter removed during a maturation process that involves nucleocytoplasmic shuttling leading to the production of a 136-nucleotide mature U8 molecule^10,11^. Termination of transcription is achieved by the integrator complex, which cleaves nascent small nuclear RNA transcripts upstream of a loosely defined 3^′^-BOX sequence^12^. We have previously shown that the 3^′^ extension of precursor human U8 forms an intramolecular duplex that masks the functionally important 5^′^ end^4^. More than 84% of patients with LCC were found to harbour at least one mutation located within this intramolecular duplex^2^, with these variants observed to increase the rate of processing of radiolabelled precursor U8 by HeLa cell nuclear extract. Mature U8 is bound by 15.5K, NOP56, NOP58 and fibrillarin at the box C/D motif, contributing to its localisation within the nucleolus, the site of pre-rRNA processing^13^. U8 also contains an LSm-binding motif that interacts with the hetero-heptameric LSm2-8 complex^14^, important in the processing and localisation of small stable RNAs within the nucleus^15^. Recently, we and others identified U8 as one of the few endogenous RNAs that form homodimers^16,17^. LCC-causative mutations are found within all of the domains outlined above^2^; however, how they affect the function of U8 remains largely uncharacterised.

RNA molecules are extensively modified, with experimental data indicating that such modifications occur at five nucleotides of U8 (RMBase v3.0 database^18^). Pseudouridylation (Ψ), the most abundant modification of cellular RNA, occurs at position n.21 located within the homodimer domain, while N^6^-methyladenosine methylation (m^6^A), the most prevalent modification of mRNA in higher eukaryotes, is detected at position n.53, adjacent to the box C motif. This m^6^A modification is present in a number of other snoRNAs, including the box C/D snoRNA U13 that also functions in pre-rRNA processing, and prevents the interaction with 15.5K, potentially modulating the timing of box C/D small nucleolar ribonucleoprotein (snoRNP) formation^19^. The remaining three modifications in U8 occur in a loop structure of the mature U8 sequence that has yet to be ascribed a function. Here, pseudouridine is detected at position n.108, and m^6^A modifications at positions n.105 and n.112. The significance of these five modifications for the function of U8 in pre-rRNA processing has not been investigated.

In this study, we used a zebrafish bioassay to assess the capacity of 50 distinct LCC-associated U8 mutations to rescue the gross morphology of the U8.3 null zebrafish embryo. By also examining the effect of these U8 mutants on 28S biogenesis and indices of translational stress, we provide independent molecular read-outs of U8 mutant functionality. We further report the effect of the LCC-causative mutations on U8 stability, identifying distinct molecular behaviours that correlate with the subdomains of U8 involved. In doing so, we reveal a post-transcriptional response that increases U8 stability when its function is hypomorphic. Finally, we show that nucleotide modifications of U8 regulate its function in 28S biogenesis, and provide evidence to suggest that such modifications play a role in disease penetrance.

## Materials and methods

### RNA extraction, cDNA synthesis and real-time quantitative PCR

RNA was isolated from homogenized n= 5 zebrafish embryos in 500 µl TRIzol (Thermo Fisher Scientific) per biological replicate according to the manufacturer’s instructions. 200 µl of the aqueous phase was taken for precipitation, with the final RNA pellet resuspended in 15 µl of RNase free water. Genomic DNA was removed using the TURBO DNA-free Kit (Thermo Fisher Scientific). Reverse transcription was performed with 1 µl of RNA (equivalent to 100-350 ng of total RNA subject to genotype) using the ProtoScript II First Strand cDNA synthesis kit (New England BioLabs) with random hexamer primers. Real-time quantitative PCR was performed using 1 µl of cDNA (80 - 90 ng when diluted 1 in 10, and 200 – 225 ng when diluted 1 in 4) with the primers listed in Extended data 1 (efficiencies ranging from 95%–105%), the SensiFAST SYBR No-ROX kit (Bioline) and a Lightcycler 480 II (Roche) or QuantStudio 5 (Thermo Fisher scientific) machine. cDNA was diluted 1 in 4 when using the Lightcycler 480 II, and 1 in 10 when using the QuantStudio5. Real-time quantitative PCR conditions were as follows: an initial denaturing step of 95°C for 10 min, followed by 40 cycles of 95°C for 30s, 60°C for 30s, 72°C for 15s, and then a dissociation step of 95°C for 15s and 60°C for 15s.

### DNA extraction, PCR for genotyping, and sequencing of an LCC cohort

Single embryos or fin clips were placed in PCR tubes with 50 µL of 50 mM NaOH and denatured for 20 min at 95°C on a Mastercycler X50 (Eppendorf). A volume of 20 µL of Tris-HCl pH 8 was added to each tube and 1 µL of the genomic DNA used for PCR amplification. When genotyping pooled embryos from 500 µl of TRIzol, the aqueous phase was removed and 150 µL of 100% ethanol was added to the organic phase, and vortexed briefly. The solution was incubated for 2-3 min at room temperature and centrifuged for 5 min at 13,300 g. The supernatant was removed, and the pellet resuspended in 50 µL of 50 mM NaOH and denatured for 20 min at 95°C on a ThermoMixer C (Eppendorf). A volume of 20 µL of Tris-HCl pH 8 was added to each tube and 1 µL of the genomic DNA used for PCR amplification. PCR amplification was performed using the primers listed in Extended data 1, Phusion polymerase (NEB), and a final concentration of 3% DMSO, employing an initial denaturing step of 98°C for 3 min, followed by 40 cycles of 98°C for 10s, 60°C for 30s and 72°C for 20-45s on a Mastercycler X50.

The *SNORD118* sequence was PCR amplified from genomic DNA from patients in our LCC cohort using primers listed in Extended data 1 that contain EcoRI and SpeI restriction enzyme sites. The PCR product was sent for Sanger sequencing (Thermo Fisher 3730xl Big Dye V3.1) at the DNA sequencing facility located within the Institute of Genetics and Cancer, University of Edinburgh. Where Sanger sequencing identified a polymorphism at position n.105 of *SNORD118*, the PCR product was column purified using a QIAquick gel extraction kit (Qiagen) according to the manufacturer’s instructions, digested with EcoRI and SpeI restriction enzymes (New England Biolabs), and subcloned into a pME vector. Sequencing was performed with M13F and M13R primers to determine which n.105 variant co-segregated with which LCC-causative mutation. The use of human material was approved by the Leeds (East) Research Ethics Committee (10/H1307/132).

### Genome editing and snoRNA transcription

Genome editing was performed as described previously^4^. Mosaic founder males harbouring U8.3 mutant alleles were crossed to female carriers of a null U8.3 mutation, and the progeny screened for phenotypes resembling null U8.3 mutants rescued by exogenous U8 transcripts i.e. embryos with slightly smaller eyes than wildtype siblings and an absence of hydrocephaly. U8.3 sequence was PCR amplified from candidate U8.3 hypomorphs and Sanger sequenced, identifying a male founder that transmitted a 47bp hypomorphic insertion allele. The male founder was then outcrossed to a wildtype female, and F1 carriers of the 47bp insertion identified as adults through genotyping of fin clips.

U8 DNA templates containing a T7 consensus sequence were PCR amplified from human or zebrafish genomic DNA (see Extended data 1 for primer sequences), and column purified using a QIAquick extraction kit (QIAGEN). Here, an additional wash with 700 µL buffer PE was performed. DNA templates were eluted in 20 µL of RNase free water. Human and zebrafish U8 snoRNAs were generated using 400ng of template DNA and a mMESSAGE mMACHINE T7 kit (Life Technologies), followed by lithium chloride precipitation overnight according to the manufacturer’s instructions, and quantitation using a NanoDrop (Thermo Fisher Scientific).

## Zebrafish U8.3 mutant rescue experiments (zebrafish bioassay)

2 nL of solution containing 500 pg of a pre-hU8 variant, phenol red and 100 pg mKate2 mRNA were microinjected into the yolk of one-cell stage zebrafish embryos through the use of the PicoSpritzer III (Parker Instruments) apparatus. Where U8 variants were found to rescue the morphology of U8.3 mutant zebrafish, genotyping was performed.

### Imaging and embryo eye size quantitation

Zebrafish embryos were anesthetized using MS-222 (Sigma Aldrich), embedded in 2.8% methyl cellulose (M0387), and imaged on an M165 FC fluorescent stereomicroscope (Leica) using an flexacam C3 (Leice) camera. The eye size of embryos at 48hpf was quantified from brightfield images using the polygon selection tool in ImageJ 1.54 to trace an outline of the left eye, with the area measured in square pixels.

Adult zebrafish were briefly anaesthetised for imaging using room temperature 1:10,000 MS-222 in PBS as per approved protocols. Fish were positioned in a petri dish, and coated with a film of solution while being imaged on a lightbox. Imaging was carried out using a Nikon DSC800 camera with 60mm lens (F36, ISO200, 2.5 second shutter speed), mounted on a Leitz Reprovit IIa overhead mount with ambient light.

### Zebrafish strains and husbandry

Embryos and adults were maintained under standard laboratory conditions. The animal experiments were reviewed by the University of Edinburgh Animal Welfare and Ethical Review Body (AWERB), and approved by the UK Animals in Science Regulation Unit (ASRU) under the Animals (Scientific Procedures) Act 1986 in strict accordance with the Home Office Code of Practice.^a,b^ Experiments were conducted following an authorised protocol endorsed by both the AWERB and the Bioresearch and Veterinary Services (BVS) department at the University of Edinburgh, under Project Licence (PP7317786) and Establishment Licence (X212DDDBD).

^a^(ASRU) AiSRU. Code of Practice for the Housing and Care of Animals Bred, Supplied or Used for Scientific Purposes. In: Office H, (ed.). UK: Williams Lea Group on behalf of the Controller of Her Majesty’s Stationery Office, 2014, p. 212.

^b^ (ASRU) AiSRU. Animals (Scientific Procedures) Act 1986. In: Office H, (ed.). United Kingdom: Williams Lea Group on behalf of the Controller of Her Majesty’s Stationery Office, 2014, p. 40.

### Bioinformatic analyses of U8 secondary structures from PARIS2 dataset

Sequencing data from the PARIS2 data set^20^ was downloaded in FASTQ format. Chimeric reads were called and annotated using hyb with bowtie2 mapping to a human transcriptome database, including an extended human U8 sequence (nucleotides 1-161 with an extension of 50nt to its 5’ and 3’ end extracted from its location on chromosome 17)^21^. U8:U8 homodimers were also identified using a method described by Gabryelska and colleagues^16^. U8 secondary structures were predicted using comradesFold^22^. Colour coding of base pairing is based on log2 of supporting chimeric reads generated through hyb.

### Statistics and reproducibility

Statistical analyses were performed using GraphPad Prism 10 or Microsoft Excel software. Raw data is summarised in Extended data 2. For all analyses, p < 0.05 was considered statistically significant. Statistical methods were not used to predetermine sample size, which varies between experiments. Experiments were not randomized. The investigators were not blinded to allocation during experiments and outcome assessment. For Figure S6 panel D, significance was determined using a Mantel-Cox test. For all other statistical analyses, significance was determined using an unpaired t test. The number of biological replicates upon which significance was determined is specified in the figure legends. Allelic variation within U8 was evaluated using pairwise two-tailed Mann-Whitney U tests, implemented via the SciPy package in Python. To account for multiple comparisons, p-values were adjusted using a false discovery rate (FDR) correction. Allele counts were obtained from gnomAD v.4.1.0, including only variants with the “PASS” quality filter.

## Results

### *In vivo* mutagenesis screening classifies LCC-associated U8 mutations as hypomorphic or null alleles

We previously reported that microinjection of wildtype human precursor *U8 (pre-hU8)* rescued the morphology of zebrafish U8.3^-/-^ embryos, thereby demonstrating evolutionarily conservation of function^4^ (Fig 1A). However, we did not assess the effect of the rescue mediated by exogenous *pre-hU8* at a molecular level. Thus, here we measured *28S* levels in U8.3^-/-^ zebrafish embryos following injection with *pre-hU8*. In doing so, we observed a partial rescue of *28S* biogenesis compared to uninjected U8.3^-/-^ embryos (Fig 1B), indicating a hypomorphic function of *hU8* in our zebrafish bioassay. We also observed significant induction of the interferon stimulated gene (ISG) *ifnphi1* in U8.3^-/-^ mutants, consistent with translational stress secondary to cGAS activation by colliding ribosomes^23^ (Fig 1C). Distinct from *28S, ifnphi1* upregulation was completely alleviated by microinjection of *pre-hU8*.

**Figure 1.**
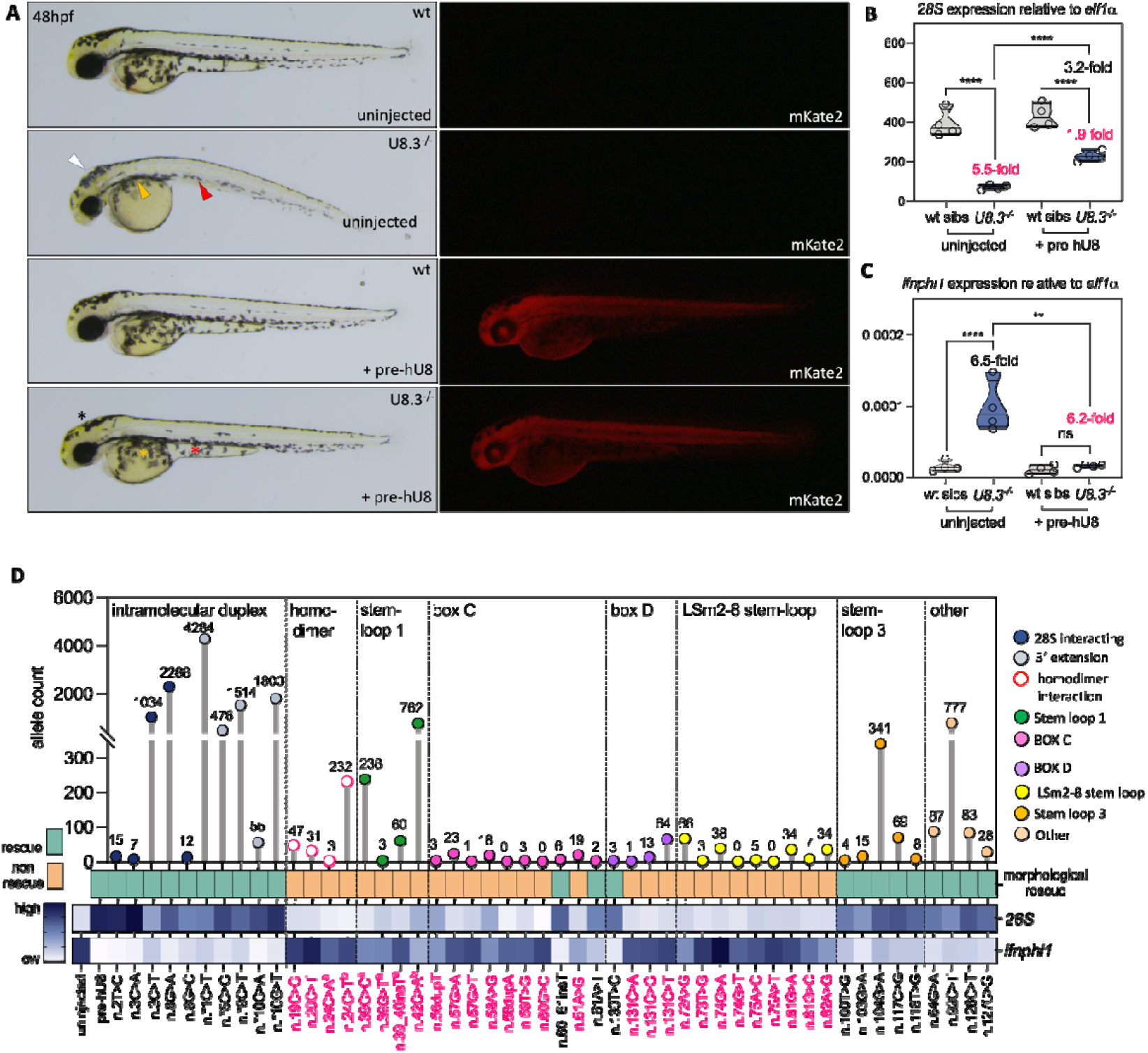
A zebrafish bioassay characterises LCC-causative U8 mutations into hypomorphic and null alleles. **A**.) Representative brightfield and fluorescent images of the indicated genotypes, exogenous snoRNA and fluorescent protein taken at 48 hours post fertilisation (hpf). White, red and orange arrowheads denote, respectively, hydrocephaly, aberrant yolk extension and impaired migration of melanocytes over the yolk. Black, red and orange asterisks denote, respectively, rescued hindbrain, yolk extension and melanocyte migration. **B**.) qRT-PCR of *28S* at 48hpf in the indicated genotypes injected with the noted exogenous snoRNA. **C**.) qRT-PCR of *ifnphi1* at 48hpf in the indicated genotypes injected with the noted exogenous U8 snoRNA. **D**.) Allele count for each LCC-associated mutation taken from gnomAD v4.1.0 (topmost panel); capacity of each LCC-associated mutation to rescue the morphology of the U8.3^-/-^ mutant embryo (middle panel). Average expression of *28S* and *ifnphi1* from U8.3^-/-^ mutants microinjected with the indicated precursor U8 mutants shown in heatmap form. Heatmap values are from qRT-PCR data normalised to *elf1a* in 3-6 biological replicates. Hypomorphic and null LCC-associated U8 mutant snoRNAs are labelled in black and red respectively. ^a^denotes U8 mutants that are detected in homozygous form in patients with LCC (n.39G > C, n.39G >T) or in control individuals annotated in the gnomAD v4.1.0 database (n.39_40insT), or co-occur with known null alleles in patients with LCC (n.24C > A co-occurs with n.82A >G), indicating hypomorphic function since homozygous null alleles are incompatible with life. ^b^denotes U8 mutants whose functionality is uncertain due to evidence that the affected nucleotides localise to regions of U8 that are not conserved at the sequence level in zebrafish. Intramolecular duplex refers to the structure that forms between the 5’ end and 3’ end of precursor human U8.

Since *SNORD118* is non-protein encoding, in silico prediction of the pathogenicity of U8 variants observed in LCC is not currently possible. Thus, we used our zebrafish bioassay to generate proof-of-pathogenicity data by assessing the function of 50 previously published LCC-associated mutations. Firstly, U8 variants were characterised according to whether or not they conferred a morphological rescue of the hydrocephaly, melanocyte migration and yolk extension defects observed in U8.3^-/-^ null embryos (Fig 1D). Representative examples of these rescue experiments are shown for mutations that localise to the intramolecular duplex important for processing of pre-hU8, the LSm2-8 binding motif, and the box C and D motifs, along with a summary of the effect of each of the 50 LCC-associated mutations on specific morphological features of the U8.3^-/-^ mutant (Fig S1A-D). Rescue, or not, of all three features was consistent for 49 of 50 variants (with the n.3C > T variant rescuing the hydrocephaly and yolk extension defects, but not the melanocyte migration) (Fig S1D), on which basis mutations were defined as either functional (providing a morphological rescue), and thus hypomorphic alleles, or non-functional (providing no morphological rescue), and thus null alleles - given their association with LCC.

Overall, we found that variants conferring a phenotypic rescue of the U8.3^-/-^ mutant were also associated with increased *28S* biogenesis and suppression of *ifnphi1*, and localised to either the intramolecular duplex required for processing of pre-U8 (an interaction detected in the PARIS2^20^ dataset, Fig S2), or stem-loop 3 of U8 (Fig 1D, Fig S1E, F). In contrast, those mutations that did not rescue the U8.3^-/-^ mutant phenotype and *28S* levels or *ifnphi1* induction, clustered within the homodimer domain, stem-loop 1, box C and D motifs, and LSm2-8 stem-loop of the mature U8 secondary structure (Fig 1D, Fig S1E, F). Concordant with these morphological and molecular data, we observed a correlation between the capacity of mutant variants to confer a phenotypic rescue and the frequency of the mutant allele in the human population (taken from the gnomAD v.4.1.0 database); with null variants observed in domains of U8 known to mediate interaction with protein (box C, box D, LSm2-8 stem-loop) being very rare, and the highest frequency of mutant alleles seen for hypomorphic variants situated in the intramolecular duplex that forms between the 5^′^ end and 3^′^ extension of precursor U8 (Fig 1D).

Rare autosomal recessive disease cohorts are typically enriched for consanguinity and molecular homozygosity. In contrast, 76 of 81 (94%) patients studied here were compound heterozygous for two mutant alleles. Such a mutational landscape suggests a threshold of U8 function, below which is incompatible with life (i.e. selection against homozygosity for a null allele), and above which LCC does not manifest (i.e. no enrichment for homozygous hypomorphic alleles in LCC cohorts). Surprisingly then, LCC patients have been recorded as homozygous for n.39G >T and n.39G > C, both of which failed to confer a rescue in our zebrafish assay. Given that these variants must retain some function in humans, we noted that stem-loop 1, which includes n.39, is poorly conserved between humans and zebrafish (Fig S3A, B), with the introduction of a G > C transversion at the only other G nucleotide (n.35) in stem-loop 1 of zebrafish pre-U8.3 also sufficient to abolish function (Fig S3C). These data indicate a zebrafish-specific intolerance of this region to mutation, which we suggest is possibly also relevant to the n.24C > A (seen to cause LCC in combination with the n.82A > G null mutation) and n.42G > A variants, both classified as null in our assay but which are seen at an unexpectedly high frequency in gnomAD v.4.1.0 (Fig 1D). An analysis of the interaction between U8 and 28S from the PARIS2 dataset indicates n.24 and stem-loop 1 mediates interaction with rRNA (Fig S4), suggesting disruption of such a duplex may underpin hypomorphic LCC-causative mutations in this region, in contrast to more common hypomorphic mutations that affect the processing of precursor U8. Taking into account these observations and the results of our zebrafish bioassay data, we classify 26 LCC-associated mutations as hypomorphic, and 22 as null (Table S1).

Relating to hypomorphic mutations, we previously reported a 60-year-old clinically asymptomatic female, compound heterozygous for two alleles (n.*1C > T, n.*9C > T) behaving as hypomorphs in our zebrafish assay, mother to two children each carrying a different maternally derived mutant allele in combination with a paternally inherited whole-gene deletion^2^ (Fig S5). This finding, together with the observation of rare individuals on gnomAD v.4.1.0 homozygous for each of these two mutations, is consistent with biallelic status for some hypomorphic mutations being insufficient to cause LCC. At the same time, 5 of 81 (6%) patients studied here were homozygous for a hypomorphic mutant allele, and a further 12 of 81 (15%) patients were compound heterozygous for two alleles classified as hypomorphic (Table S2). These data suggest either variability in the function of variants classified here as hypomorphic and which our zebrafish model is not sensitive enough to identify, or the effect of environmental or other genetic factors in determining clinical outcome.

### The stability of U8 snoRNA increases through a post-transcriptional mechanism when its function is impaired

As part of our analysis of the molecular consequences of LCC-associated mutations, we quantified exogenous wildtype *hU8* following injection of *pre-hU8* into the zebrafish embryos used in our bioassay. In doing so, we observed increased levels of *hU8* in U8.3^-/-^ embryos compared to wildtype siblings, despite the introduction of equal amounts of *pre-hU8* (Fig 2A). To determine if a similar effect was seen with endogenously expressed genes, we generated a novel zebrafish U8.3 hypomorphic mutant containing a 47bp insertion (U8.3^47ins^) that triplicates regions of the homodimer domain and 28S interacting nucleotides of U8 (Fig 2B). Importantly, the expression of the endogenous *zU8*.*3*^47ins^ allele was increased in U8.3^47ins/-^ embryos where *28S* biogenesis is compromised, compared to their U8.3^47ins/+^ siblings (Fig 2C-E). Of note, U8.3^47ins/-^ larvae exhibit embryonic lethality, whereas U8.3^47ins/47ins^ larvae survive to adulthood, indicating that U8.3 is subject to a gene dosage effect (Fig S6), consistent with a close to 2-fold reduction of zU8.3 in U8.3^+/-^ embryos compared to U8.3^+/+^ embryos (Fig 2C).

**Figure 2.**
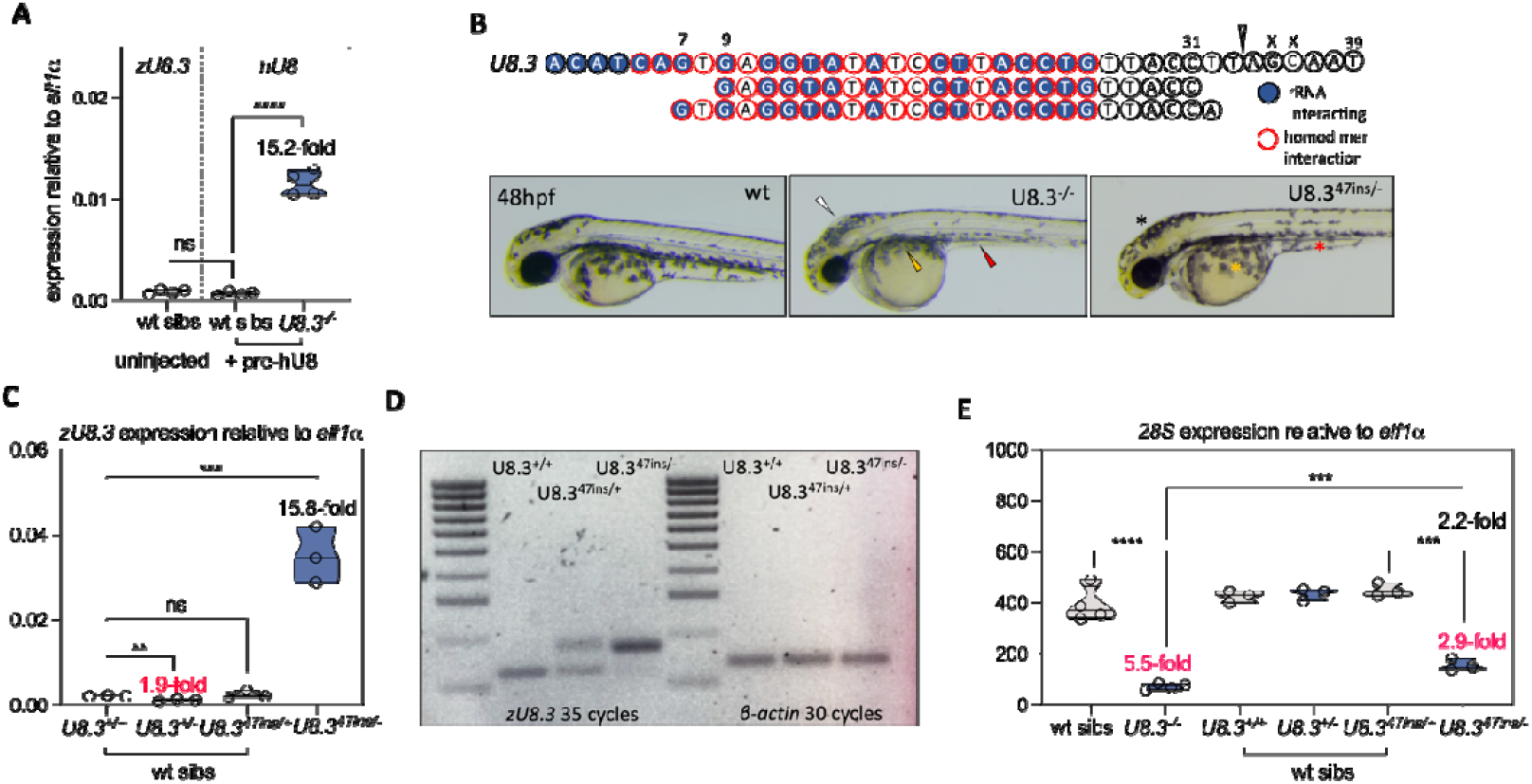
U8 increases stability through a post-transcriptional mechanism when hypomorphic in function. **A**.) qRT-PCR of endogenous *zU8*.*3* and exogenous *hU8* snoRNAs at 48hpf in the indicated genotypes. *zU8*.*3* and exogenous *hU8* are expressed at equal levels in wildtype embryos. **B**.) Schematic depicting the 47-base pair (bp) insertion that results in a hypomorphic allele of zU8.3. This allele comprises insertion of the second row of sequence, followed by the third row of sequence (49bp in total), at the position indicated by the white arrowhead in the top row of sequence, which is the first 39bp of wildtype zU8.3. Nucleotides 35 and 36 are deleted from the hypomorphic allele (represented by crosses). Below the schematic are representative brightfield images of a wildtype (wt) zebrafish embryo and indicated genotypes. U8.3^-/-^ embryos exhibit hydrocephaly (white arrowhead), reduced migration of melanocytes over the yolk (orange arrowhead) and reduced development of the yolk extension (red arrowhead). These features are rescued in the U8.3^47ins/-^ hypomorph, denoted by the black, orange and red asterisks respectively. **C**.) qRT-PCR of *U8*.*3* at 48hpf in the indicated genotypes. **D**.) RT-PCR of *zU8*.*3* and *β-actin* at 48hpf in the indicated genotypes. **E**.) Quantitative RT-PCR (qRT-PCR) of *28S* at 48hpf in the indicated genotypes.

Summarising, in two distinct systems, we recorded increased expression of either exogenous *hU8* or endogenous *zU8*.*3* when U8 function is compromised. These data suggest the activation of an evolutionarily conserved post-transcriptional mechanism that we hypothesise to represent a compensatory physiological response to reduced U8 activity.

### The effect of LCC-associated mutations on U8 stability varies according to U8 domain

In our zebrafish models, the stability of injected exogenous *hU8* (Fig 2A) and endogenous *zU8*.*3* (Fig 2C) differed according to background genotype, and whether U8 function is impaired. Next then, we assessed the effect of LCC-causative mutations on U8 stability by systematically quantifying the levels of *hU8* following injection of mutant *pre-hU8* variants in both wildtype sibling and U8.3^-/-^ embryos. Mutations previously determined as diverging in function in the zebrafish bioassay (i.e. n.24C > T, n.24C > A and the stem-loop 1 mutations involving n.39 and n.42: see above) were excluded from this analysis. In this way, we noted distinct molecular behaviours according to the localisation of mutant nucleotides within the different domains of U8.

Firstly, considering hypomorphic variants conferring a rescue of the U8.3^-/-^ phenotype, we observed that mutations within the intramolecular duplex required for processing of precursor U8 [assessed previously using Hela nuclear extract processing assays^1,4^, and confirmed here for three previously unstudied variants (Fig S7)], exhibited similar stability to wildtype *hU8* in wildtype or U8.3^-/-^ mutant embryos (Fig 3). The same was true for the n.92C

**Figure 3.**
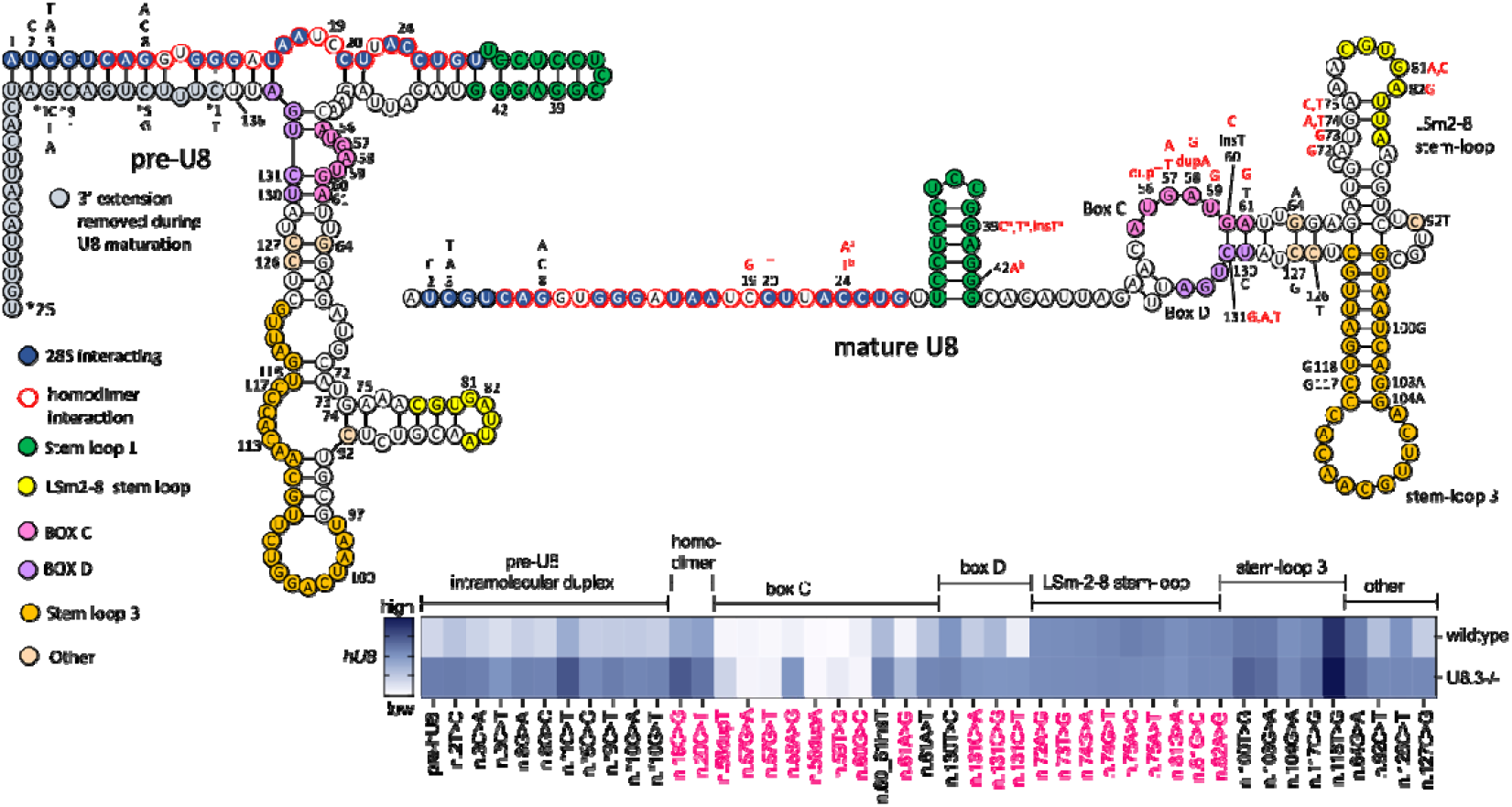
The effect of LCC-causative mutations on U8 stability clusters according to U8 structural and protein interacting domains. Schematic depicting the secondary structure of precursor and mature human U8 snoRNA after removal of the 3^′^ extension. Average expression of exogenous *hU8* from wildtype and U8.3^-/-^ embryos microinjected with the indicated precursor U8 variants is shown in heatmap form. Heatmap values are from qRT-PCR data normalised to *elf1a* in 3-6 biological replicates. Hypomorphic LCC-associated U8 mutant snoRNAs are labelled in black, and null alleles labelled in red. ^a^denotes U8 mutants that are detected in the homozygous state in patients with LCC (n.39G>C, n.39G >T), or in the gnomAD v4.1.0 database (n.39_40insT), or co-occur with known null alleles in patients with LCC (n.24C > A co-occurs with n.82A >G), indicating hypomorphic function since homozygous null alleles are incompatible with life. ^b^denotes U8 mutants whose functionality is uncertain due to evidence that the affected nucleotides localise to regions of U8 that are not conserved at the sequence level in zebrafish. The effect on U8 stability of mutations denoted ^a^ or ^b^ is not included due to this lack of conservation.

T and n.127C > G variants, and the box C mutations n.60_61insT and n.61A > T (Fig 3, Fig S8). In contrast, stem-loop 3 hypomorphic mutations displayed significantly increased stability in wildtype embryos, which was equivalent to the expression observed in U8.3^-/-^ embryos (Fig 3, Fig S8). The box D mutation n.130T > C, and n.64G > A and n.126C >T, also displayed significantly increased stability in wildtype embryos, but with comparatively higher expression in U8.3^-/-^ embryos (Fig 3, Fig S8). Secondly, when considering null alleles, again in comparison to wildtype pre-hU8, the injected homodimer mutants n.19C > G and n.20C > T displayed increased stability in wildtype embryos, and even greater stability in U8.3^-/-^ embryos. In contrast, the majority of box C mutations were highly destabilising in both genotypes (Fig 3, Fig S8). Null mutations involving the LSm2-8 stem-loop exhibited elevated stability in wildtype embryos, equivalent to the expression observed in U8.3^-/-^ embryos (a behaviour similar to hypomorphic mutations involving stem-loop 3) (Fig 3, Fig S8). Finally, the box D null alleles involving n.131C displayed broadly comparable stability to wildtype *hU8* in wildtype embryos, but significantly reduced stability compared to wildtype *hU8* on a U8.3^-/-^ genetic background (Fig 3, Fig S8).

Taken together, the behaviour of LCC-associated variants in our system varied according to functional severity (hypomorph versus null), zebrafish genetic background (wildtype or U8.3^-/-^ mutant), and the domain of U8 involved. As such, the molecular mechanisms that underpin the pathogenicity of the LCC-associated mutations differ between domains of U8, providing insight into the role of these domains in U8 and wider snoRNA biology.

### Pseudouridine (Ψ) and m^6^A modified nucleotides impact U8 function and may play a role in disease penetrance

The mechanism underpinning the increase in U8 stability when its function is hypomorphic must be post-transcriptional, as this response is observed with exogenously introduced U8 transcripts. Examination of RMBase 3.0 indicates that human U8 is post-transcriptionally modified at five nucleotides (Fig S9A), with Ψ (n.21, n.108) and m^6^A (n.53, n.105, n.112) modified nucleotides lying within PUS4-associated^24^ (Fig S9B) and 5^′^-DRACH-3^′^ (Fig S9C) motifs, respectively. Of note, three (n.105, n.108, n.112) of these modified nucleotides are situated in the evolutionarily conserved 11-nucleotide loop region of stem-loop 3 (Fig S9A, S9D).

To explore the possibility that Ψ or m^6^A modification might impact U8 behaviour in zebrafish, we mutated each of the five modifiable nucleotides to prevent their post-transcriptional modification, and then assessed the effect on U8 in our bioassay. The choice of mutation was informed by the allelic frequency observed for each of the modified nucleotides in the human population according to gnomAD v4.1.0, selecting the highest frequency variants on the assumption that they are least likely to disrupt U8 function through alternative mechanisms, such as by altering the secondary structure of U8.

Considering, firstly, Ψ modified nucleotides, we found that mutation of n.21T abrogated the ability of U8 to rescue the morphology of the U8.3^-/-^ mutant (Fig 4A). At the molecular level, n.21T > G phenocopied the null n.19C > G and n.20C > T homodimer LCC-causative mutations in demonstrating a significant increase in stability in U8.3^-/-^ mutant embryos in the absence of a morphological rescue (Fig 4B, C, D). In contrast, and unexpectedly, mutation of n.108T to n.108G was associated with enhanced melanocyte migration and a larger eye size compared to the morphological rescue achieved by microinjection of wildtype *pre-hU8* (Fig 4A). Controlling for developmental variability between embryos collected from pooled groups of genetically distinct zebrafish parents, we observed the same enhanced melanocyte migration and increased eye size with mutant (n.108T > G) versus wildtype *pre-hU8* injected into sibling embryos derived from the same parents (Fig S10A, B). Further consistent with these observations, the lower jaw angle of zebrafish embryos was less developmentally delayed in U8.3^-/-^ mutants injected with n.108T > G compared to *pre-hU8* (Fig S10A). At a molecular level, n.108T > G exhibited equal stability in both wildtype and U8.3^-/-^ mutant embryos (Fig 4B), rescued *28S* biogenesis (Fig 4C) and suppressed *ifnphi1* (Fig 4D). Of note, n.108T > G did not significantly increase *28S* biogenesis beyond that of wildtype U8 (Fig 4C), and while inactivation of tp53 confers a partial rescue of the U8.3^-/-^ mutant^4^, n.108T> G did not suppress expression of the tp53 target gene *mdm2* from levels seen in uninjected U8.3^-/-^ mutants or U8.3^-/-^ mutants injected with wildtype *pre-hU8* (Fig 4E). Taken together, these data indicate that n.108T confers a negative regulatory activity on U8 function, which may be dependent on the presence of a pseudouridine modification; although it cannot be determined whether pseudouridylation positively or negatively regulates n.108T function from this experiment. The almost complete absence of allelic variation at n.21 and n.108 of U8 in humans suggests that these nucleotides, and their associated modifications, play an important role in U8 mediated homeostasis (Fig S11).

**Figure 4.**
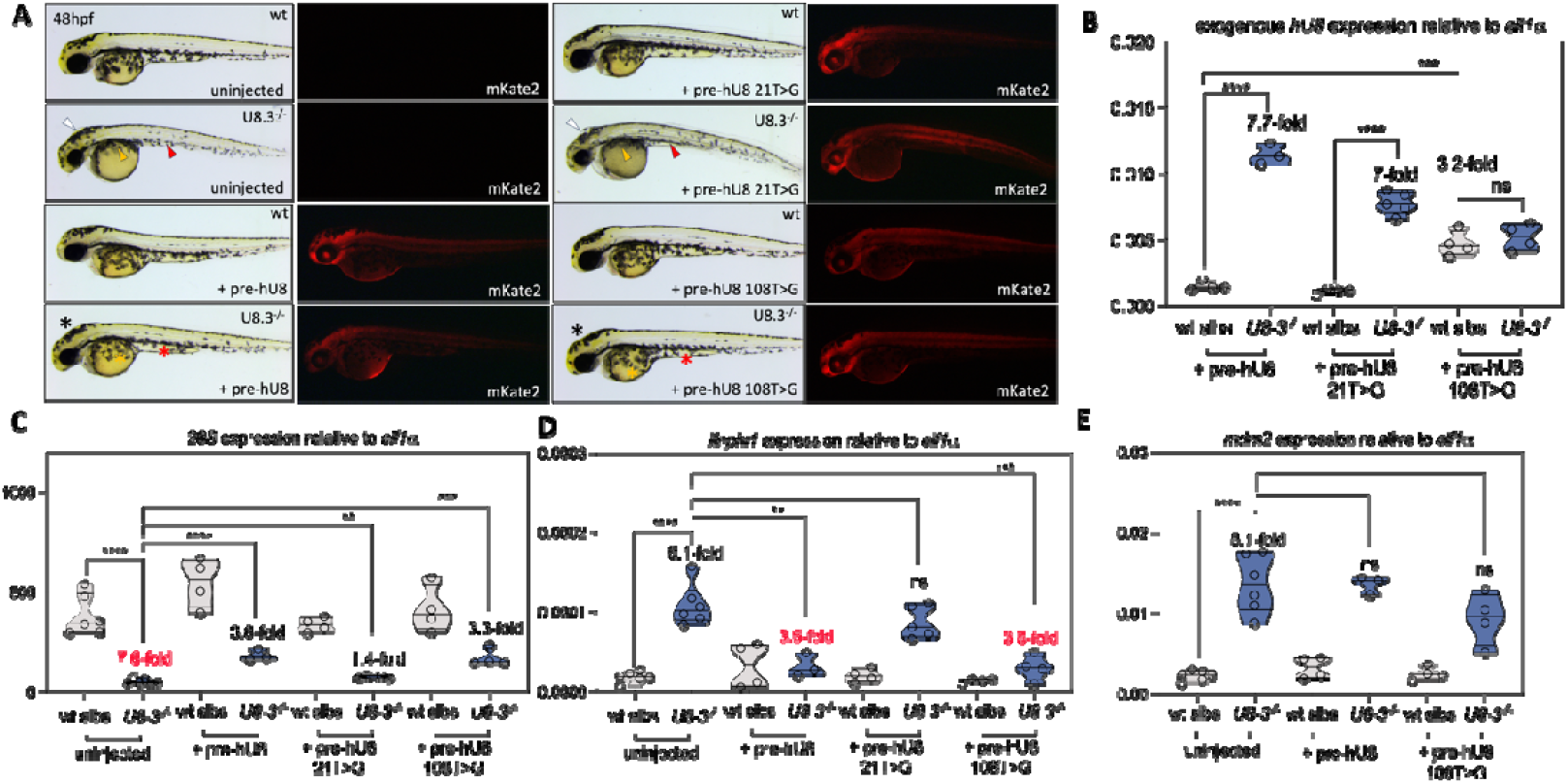
Pseudouridine modified nucleotides positively and negatively regulate U8 function. **A**.) Representative brightfield and fluorescent images of the indicated genotypes, exogenous snoRNAs and fluorescent protein is given at 48hpf. **B**.) qRT-PCR of the indicated exogenous U8 snoRNAs at 48hpf in the given genotypes. **C**.) qRT-PCR of *28S* for the indicated snoRNAs at 48hpf in the given genotypes. **D**.) qRT-PCR of *ifnphi1* for the indicated snoRNAs at 48hpf in the given genotypes. **E**.) qRT-PCR of *mdm2* for the indicated snoRNAs at 48hpf in the given genotypes. White, red and orange arrowheads denote, respectively, hydrocephaly, aberrant yolk extension and impaired migration of melanocytes over the yolk. Black, red and orange asterisks denote, respectively, rescued hindbrain, yolk extension and melanocyte migration. pre - precursor. Significance determined by unpaired t-test. Each biological replicate is indicated by a circle. Red fold change indicates decrease, black fold change indicates increase.

Next, we examined the functional requirement for the m^6^A modified nucleotides in our zebrafish bioassay. n.53A > G, n.105A > T and n.112A > G were selected to block m^6^A modification, noting that n.53A > G substitutes an A:U for a G:U interaction, thereby likely maintaining U8 secondary structure, and that n.105T is the predominant allele in the human population. While all three mutants were able to rescue U8.3^-/-^ morphology (Fig 5A), *28S* and *ifnphi1* expression (Fig 5B, C), their impact on U8 stability in U8.3^-/-^ embryos was distinct from that of wildtype *hU8* (Fig 5D), consistent with a functional role of m^6^A modification. Notably, 50 of 53 patients with LCC in our cohort (94%) harbour the n.105T variant on an allele assessed as hypomorphic in this study (Table S3), indicating that almost the entire LCC cohort cannot m^6^A modify functional U8 at this nucleotide, compared to only 40% of the general population. Of the three exceptions that we recorded, while F344 is homozygous for the n.105A variant, this patient is also homozygous for another stem-loop 3 variant, n.113C > T (Table S3); and in F2089, the n.105A allele co-segregates with a hypomorphic n.*9C > T mutation and another rare variant within the stem-loop 3 structure (n.119G > T) (Table S3). It may be then that n.113C > T and n.119G > T sufficiently compromise the function of stem-loop 3 in LCC patients to phenocopy the effect of n.105T in the rest of our cohort.

**Figure 5.**
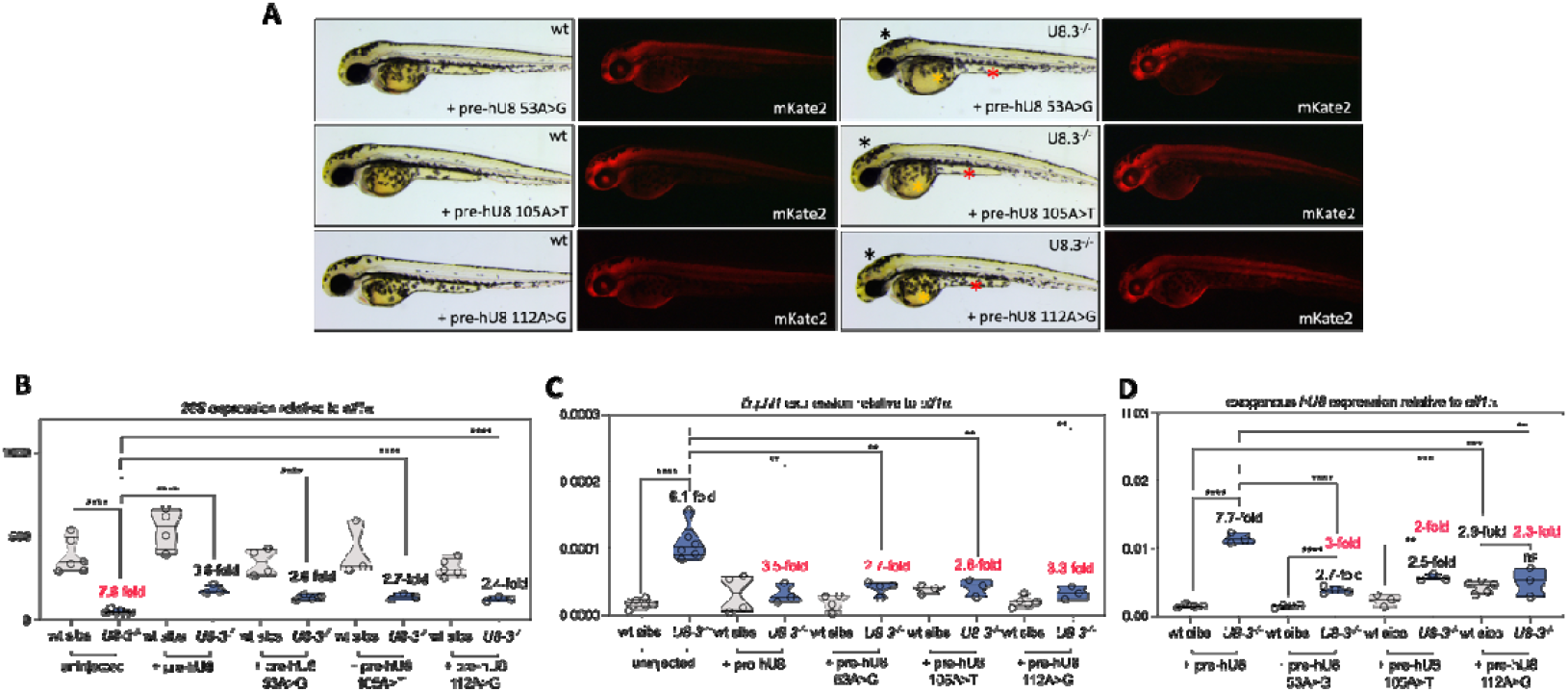
Mutation of m^6^A modified nucleotides alters the stability of U8. **A**.) Representative brightfield and fluorescent images of the indicated genotypes, exogenous snoRNAs and fluorescent protein is given at 48hpf. **B**.) qRT-PCR of *28S* for the indicated snoRNAs at 48hpf in the given genotypes. **C**.) qRT-PCR of *ifnphi1* for the indicated snoRNAs at 48hpf in the given genotypes. **D**.) qRT-PCR of the indicated exogenous U8 small nucleolar RNAs (snoRNAs) at 48hpf in the given genotypes. White, red and orange arrowheads denote, respectively, hydrocephaly, aberrant yolk extension and impaired migration of melanocytes over the yolk. Black, red and orange asterisks denote, respectively, rescued hindbrain, yolk extension and melanocyte migration. pre - precursor. Significance determined by unpaired t-test. Each biological replicate is indicated by a circle. Red fold change indicates decrease, black fold change indicates increase.

### Stem-loop 3 of U8 is essential for 28S biogenesis

Several lines of evidence suggest a significant role for stem-loop 3 in U8 biology, specifically: 1) the loop sequence is conserved across evolution (Fig S9D); 2) the 11-nucleotide loop sequence contains three (n.105, n.108, n.112) of the five modifiable nucleotides of U8 (Fig S9A); 3) five LCC-causative mutations localise to the stem structure of stem-loop (Fig 3); 4) a patient from our cohort (F344) is homozygous for both a n.8G > C and n.113C > T variant, with the n.113 nucleotide part of the loop structure of stem-loop 3. This latter individual is notable for exhibiting the earliest recorded age at disease onset of any patient in our LCC cohort, leading us to hypothesise that the n.113C > T variant contributes to pathology in addition to the hypomorphic n.8G > C mutation previously shown to affect pre-U8 processing^4^.

While null mutations cluster in known protein binding domains characterised by minimal allelic variation, nucleotides constituting the loop sequence of stem-loop 3 exhibit allelic variation more comparable to regions of U8 known to mediate interactions with rRNA i.e. the intramolecular duplex and stem-loop 1 (Fig 1D, Fig S11, Table S4). Noting that LCC-causative mutations in the intramolecular duplex that forms between the 3^′^ extension of pre-U8 and its 5^′^ end preferentially involve G:C interactions (Fig 1D), we hypothesised that the loop sequence of stem-loop 3 might also mediate RNA-RNA interactions. To investigate this possibility, we mutated the n.110C and n.113C nucleotides that localise to this structure. In doing so, we found that C > T transition of n.110 and n.113 was severely deleterious at both a morphological and molecular level, with combined mutations at these sites sufficient to render U8 null (Fig 6), and abrogate the increase in the expression of wildtype *hU8* seen in injected U8.3^-/-^ embryos (Fig 6B). These data indicate that the loop sequence of stem-loop 3 plays an essential role in pre-rRNA processing.

**Figure 6.**
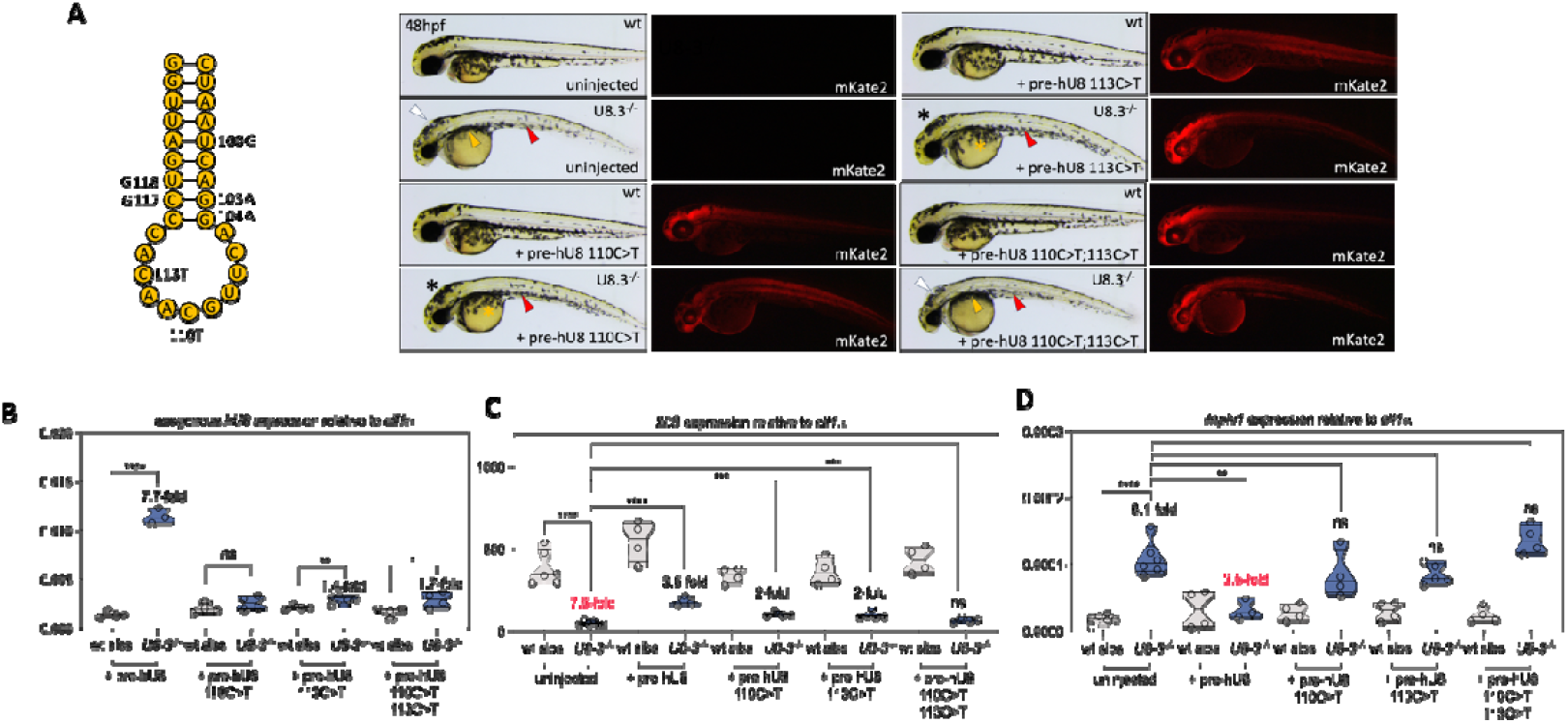
The loop sequence of stem-loop 3 is critical for the function of U8 in 28S biogenesis. **A**.) Representative brightfield and fluorescent images of the indicated genotypes, exogenous snoRNAs and fluorescent protein is given at 48hpf. **B**.) qRT-PCR of the indicated exogenous U8 snoRNAs at 48hpf in the given genotypes. **C**.) qRT-PCR of *28S* for the indicated snoRNAs at 48hpf in the given genotypes. **D**.) qRT-PCR of *ifnphi1* for the indicated snoRNAs at 48hpf in the given genotypes. White, red and orange arrowheads denote, respectively, hydrocephaly, aberrant yolk extension and impaired migration of melanocytes over the yolk. Black, red and orange asterisks denote, respectively, rescued hindbrain, yolk extension and melanocyte migration. pre - precursor. Significance determined by unpaired t-test. Each biological replicate is indicated by a circle. Red fold change indicates decrease, black fold change indicates increase.

## Discussion

Here we report the functional and molecular assessment of 50 LCC-associated mutations in *SNORD118*, encoding the vertebrate specific box C/D U8 snoRNA. We also provide a targeted investigation of five Ψ and m^6^A modified nucleotides, and three selected nucleotides in stem-loop 3 of U8. In doing so, these studies represent an important contribution to defining the molecular architecture and biological functions of U8.

Cohorts of individuals affected by rare autosomal recessive disorders are typically enriched for parental consanguinity and allelic homozygosity. Strikingly then, consanguinity was noted in only three of 56 families in a large cohort of individuals with LCC that we previously reported^2^, and 94% of the patients studied here were compound heterozygous for two distinct mutations in *SNORD118*. Such a mutational landscape suggests a threshold of U8 function, below which is incompatible with life (i.e. selection against homozygosity for a null allele), and above which LCC does not manifest (i.e. no enrichment for homozygosity for hypomorphic alleles in LCC cohorts). Indeed, our morphological rescue data support the hypothesis that LCC typically results due to the combination of a hypomorphic (functional) and null (non-functional) mutation, with this gene dosage effect evolutionarily conserved to zebrafish (where a single U8.3^47ins^ allele in combination with a null allele is embryonic lethal, while two copies of the hypomorphic U8.3^47ins^ allele are adult viable). Overall, we found that variants conferring a morphological rescue of the U8.3^-/-^ zebrafish mutant were also associated with increased 28S biogenesis and suppression of *ifnphi1* induction. Further, we note a correlation between the capacity of mutant variants to confer a phenotypic rescue and the frequency of the mutant allele in the human population, suggesting that nucleotide allelic frequency can serve as a useful metric in assessing the molecular pathogenicity of a given U8 variant.

As part of the characterisation of U8 function using our bioassay, we recorded increased stability of exogenous *hU8* and endogenous *zU8*.*3* when U8 function is hypomorphic. We interpret these data to suggest the activation of an evolutionarily conserved post-transcriptional mechanism as a physiological response to reduced U8 activity. We also observed the clustering of null and hypomorphic mutations in, respectively, domains of U8 mediating interaction with protein (i.e. box C, box D, LSm2-8 stem-loop) or the intramolecular duplex (that forms between the 5’ end and 3’ extension of precursor U8) / stem-loop 3. Again, as for mutation type and allelic frequency, mutations displaying similar effects on stability clustered according to U8 domain, providing further evidence of shared domain-specific pathogenic mechanisms, which likely hold significance for basic U8 and snoRNA biology. Notably, U8 mutant stability did not necessarily correlate with the hypomorphic or null status of individual mutant alleles. Standard approaches for characterising the function of non-coding RNAs typically involve either their knockdown or knockout, lacking the capacity to identify the molecular characteristics captured in this study. Likewise, although saturated mutagenesis screens using cellular proliferation as a readout have the potential to differentiate hypomorphic from null mutations in non-coding RNAs, this approach cannot yet assess effects on RNA stability^25^. As such, the *in vivo* zebrafish system that we describe is unique in providing a relatively high throughput means to investigate the pathogenicity and molecular behaviour of mutant non-coding RNA alleles.

Minimal free energy (MFE) calculations assessing RNA folding efficiency can be used to infer the effect of sequence variants in non-protein encoding RNAs. Using psoralen analysis of RNA interactions and structures 2 (PARIS2), Zhang et al. showed that U8 interacts with U13 and can take up alternative conformations during rRNA processing^20^. In this study, the authors also considered MFE changes induced by 32 LCC-associated mutations across ten RNA duplexes, with the results diverging in important ways from the data obtained using our bioassay. Thus, firstly, while Zhang et al. found C box mutations to be neutral or stabilising, we observed these variants to be the most destabilising of the 50 LCC-causative mutations that we assessed. Secondly, Zhang et al. suggested mutations in the homodimer domain to be highly disruptive. In contrast, while such mutations can disturb the palindromic homodimer interaction, we found the n.19C > G and n.20C > T U8 mutants to be more stable than wildtype U8, indicating that U8 monomers are themselves relatively stable.

RNA modifications are widespread, reversible and associated with fundamental biological processes^26^. In U8, m^6^A modification of n.53A has been proposed to regulate box C/D snoRNP assembly by impairing interaction with 15.5K^19^. Notably, the consensus site for this modification is poorly conserved between humans, mouse and some zebrafish U8 paralogues (Fig S12), and we found that mutation of any of the three m^6^A modified nucleotides in human U8 did not significantly impair the ability of U8 to rescue the morphology of the U8.3 null mutant zebrafish. Nevertheless, we did identify an effect of these m^6^A modified nucleotides on U8 stability when its function is hypomorphic. Perhaps relevant to this latter observation, m^6^A modification can trigger mRNA degradation through UPF1-dependent decapping^27^, with the U8-specific binding partner Nudt16 also possibly acting through m^6^A-dependent decapping^28,29^. Several recently developed methodologies may facilitate clarification of the role of RNA modification in U8 biology in future studies; in particular, nanopore sequencing, which can define RNA m^6^A and Ψ modifications with single nucleotide resolution in a quantitative manner^30–32^, and TREX^33^, which allows for site-specific mapping of protein binding partners to RNA.

Leveraging the power of human genetics and an *in vivo* zebrafish bioassay, our study highlights a previously unappreciated role of the loop sequence of U8 stem-loop 3 in pre-rRNA processing. While comparatively increased levels of human allelic variation within the loop 3 sequence of U8 might suggest an interaction with rRNA, we could not detect such an interaction in our analysis of a PARIS2 dataset (data not shown). It may be that this structure interacts with pre-rRNA transiently, and/or in a dynamic manner, possibly mediated by the encompassed three modified nucleotides (n.105, n.108, and n.112). Consistent with this hypothesis, several conformations of U8 were detected by PARIS2 in human cells^20^. Assessing the effect of the role of modified nucleotides in the stem-loop 3 loop sequence in putative RNA interactions is challenging, since mutagenesis aimed at abrogating these modifications may also affect interactions within an RNA duplex. Thus, although we observed mutation of modified nucleotides in stem-loop 3 to affect U8 stability, we cannot attribute this effect solely to the function of the modification per se. Furthermore, while mismatches can be tolerated in regions that interact with rRNA^25^, species-specific differences resulted in certain mutations that must be functional in humans (particularly, those involving n.24 and n.39) to exhibit no or minimal activity in our system.

Ribosomopathies typically involve the haematopoietic system or craniofacial and skeletal structures derived from the neural crest^34^. In this respect, the neurologically restricted phenotype of LCC is unusual. Given that RNA modifications are particularly prevalent in the brain^35,36^, it is of possible interest that U8 is both Ψ and m^6^A modified. With three exceptions, our entire LCC cohort is unable to m^6^A modify U8 at n.105 on the background of a hypomorphic allele, which we hypothesise might explain a preferential impact of U8 dysfunction in the central nervous system. Further, our finding that modified nucleotides can act as either positive or negative regulators of U8 function suggests both that post-transcriptional modifications may influence disease onset and severity (with the capacity to methylate n.105A on a functional U8 allele protective against the development of LCC), and that therapeutic approaches to modulate such modifications might confer clinical benefit.

## Supporting information

Table of oligonucleotides

Raw data used for figures

## Contributions

Zebrafish experiments: A.P.B, K.B, C.W; Processing assays: R.T.O; Bioinformatics and statistics: A.P.B, J.Y.L; Sanger sequencing: A.P.B; C.P; Experimental design: A.P.B; Conceptualisation: A.P.B. Y.J.C, G.K; Supervision: A.P.B, Y.J.C, G.K; Drafting of manuscript: A.P.B, Y.J.C, G.K, R.T.O, revision by all authors; Acquisition of funds: Y.J.C, G.K, R.T.O

## Acknowledgments

We would like to thank the staff from the MRC HGU Zebrafish facility, Advanced Imaging Resource facility and DNA sequencing facility.

We would like to thank all of the families and clinicians that have contributed to this work over the last 20 years. We would like to particularly thank Mandy Vine and her family for their unwavering encouragement.

Y.J.C acknowledges UK Medical Research Council (MRC) support including Grant Ref: MR/V000195/1Y.J.C. and an MRC Human Genetics Unit core grant (MC_UU_00035/11).

