## Supplementary material for "Human Mendelian disease and *in vivo* mutagenesis screening define the molecular architecture of the U8 snoRNA": Table of oligonucleotides

| **Application** | **Oligonucleotide sequence 5' to 3'** |
| --- | --- |
| Zebrafish U8.3 qPCR and RT-PCR | **S** GAGGTATATCCTTACCTGTTAC  **AS** GATTCGTAAAGGGGTTGCAG |
| Zebrafish U8.3 47bp allele qPCR | **S** GGTTACAGATCATGATGATTGG  **AS** GATTCGTAAAGGGGTTGCAG |
| Human U8 qPCR to wildtype U8 and LCC variants | **S** GTGGGATAATCCTTACCTG  **AS** ATCAGGGTGTTGCAAGTCCTG |
| n.19C > G qPCR | **S**  GTGGGATAATGCTTACCTG  **AS** use oligonucleotide from human U8 |
| n.20C > T qPCR | **S**  GTGGGATAATCtTTACCTG  **AS** use oligonucleotide from human U8 |
| n.24C > A qPCR | **S**  GTGGGATAATCCTTAtCTG  **AS** use oligonucleotide from human U8 |
| n.24C > T qPCR | **S**  GTGGGATAATCCTTAaCTG  **AS** use oligonucleotide from human U8 |
| n.103G > A qPCR | **S**  use oligonucleotide from human U8  **AS** ATCAGGGTGTTGCAAGTCTTG |
| n.104G > A qPCR | **S**  use oligonucleotide from human U8  **AS** ATCAGGGTGTTGCAAGTTCTG |
| n.117C > G qPCR | **S**  use oligonucleotide from human U8  **AS** ATCACGGTGTTGCAAGTCCTG |
| n.118T > G qPCR | **S**  use oligonucleotide from human U8  **AS** ATCCGGGTGTTGCAAGTCCTG |
| Human U8 qPCR to loop 3 U8 mutants | **S**  use oligonucleotide from human U8  **AS** ATCAGACAGGAGCAATCAGG |
| n.21T > G qPCR | **S** GTGGGATAATCCGTACCTG  **AS** use sequence from loop 3 qPCR |
| Zebrafish 28S qPCR | **S** ACCGTCGTGAGACAGGTTAG  **AS** TCCCACAGATGGTAGCTTCG |
| Zebrafish ifnphi1 qPCR | **S** CACACAAGGAGTCCTACGAG  **AS** GCGATGATGTCCATCCTCTG |
| Zebrafish mdm2 | PMID: 32359472 |
| Zebrafish Δ113p53 | PMID: 32359472 |
| Zebrafish elf1a qPCR | PMID: 32359472 |
| Zebrafish beta-actin2 RT-PCR | **S** GGGAAAAGATGACACAGATC  **AS** GTGACACCATCACCAGAGTC |
| Zebrafish U8.3 null genotyping | PMID: 32359472 |
| Zebrafish U8.3 47bp allele genotyping | **S** TAAACGATCGTCTCGTCCAC  **AS** GGAATGGAGTCACAGACTTAC |
| mKate2 qPCR | **S**  ACACCTGATCTGCAACCTG  **AS** GCTCGACGTATGTCTCTTG |
| Human U8 sequencing and subcloning | **S** GACAGAGAATTCCGTAACTGATCGGAGCATTC  **AS** GACAGAACTAGTCCATGTGCACAAGACTGCAG |
| DNA template for production of zebrafish mature U8.3 snoRNA | PMID: 32359472 |
| DNA template for production of zebrafish pre-U8.3 | PMID: 32359472 |
| DNA template for production of human mature U8 snoRNA | PMID: 32359472 |
| DNA template for production of human pre-U8 | PMID: 32359472 |
| DNA template for production of human pre-U8 n.2T > C | PMID: 32359472 |
| DNA template for production of human pre-U8 n.3C > A | **S**  TAATACGACTCACTATAGGGGATAGTCAGGTGGGATAATCCTTACCTG  **AS** use oligonucleotide from human pre-U8 |
| DNA template for production of human pre-U8 n.3C > T | PMID: 32359472 |
| DNA template for production of human pre-U8 n.8G > A | PMID: 35332098 |
| DNA template for production of human pre-U8 n.8G > C | PMID: 32359472 |
| DNA template for production of human pre-U8 n.19C > G | PMID: 35332098 |
| DNA template for production of human pre-U8 n.20C > T | PMID: 35332098 |
| DNA template for production of human pre-U8 n.24C > A | **S**  TAATACGACTCACTATAGGGGATCGTCAGGTGGGATAATCCTTAA  CTGTTCCTCCTC  **AS** use oligonucleotide from human pre-U8 |
| DNA template for production of human pre-U8 n.24C > T | PMID: 35332098 |
| DNA template for production of human pre-U8 n.39G > C | **S**  TAATACGACTCACTATAGGGGATCGTCAGGTGGGATAATCCT  TACCTGTTCCTCCTCCGCAGGGCAGATTAG  **AS** use oligonucleotide from human pre-U8 |
| DNA template for production of human pre-U8 n.39G > T | **S**  TAATACGACTCACTATAGGGGATCGTCAGGTGGGATAATCCT  TACCTGTTCCTCCTCCGTAGGGCAGATTAG  **AS** use oligonucleotide from human pre-U8 |
| DNA template for production of human pre-U8 n.39_40insT | **S**  TAATACGACTCACTATAGGGGATCGTCAGGTGGGATAATCCT  TACCTGTTCCTCCTCCGGTAGGGCAGATTAG  **AS** use oligonucleotide from human pre-U8 |
| DNA template for production of human pre-U8 n.42G > A | **S**  TAATACGACTCACTATAGGGGATCGTCAGGTGGGATAATCCTTACCTG  TTCCTCCTCCGGAGAGCAGATTAGAAC  **AS** use oligonucleotide from human pre-U8 |
| DNA template for production of human pre-U8 n.56dup | **S** TAATACGACTCACTATAGGGGATCGTCAGGTGGGATAATCCTTACCTGTTCCT CCTCCGGAGGGCAGATTAGAACATTGATGATTGGAGATGCATG  **AS** use oligonucleotide from human pre-U8 |
| DNA template for production of human pre-U8 n.57G > A | PMID: 32359472 |
| DNA template for production of human pre-U8 n.57G > T | **S**  TAATACGACTCACTATAGGGGATCGTCAGGTGGGATAATCCTTACCTGTTCCTC  CTCCGGAGGGCAGATTAGAACATTATGATTGGAGATGCATG  **AS** use oligonucleotide from human pre-U8 |
| DNA template for production of human pre-U8 n.58A > G | PMID: 32359472 |
| DNA template for production of human pre-U8 n.58dup | **S**  TAATACGACTCACTATAGGGGATCGTCAGGTGGGATAATCCTTACCTGTTCCTCCT  CCGGAGGGCAGATTAGAACATGAATGATTGGAGATGCATG  **AS** use oligonucleotide from human pre-U8 |
| DNA template for production of human pre-U8 n.59T > G | **S**  TAATACGACTCACTATAGGGGATCGTCAGGTGGGATAATCCTTACCTGTTCCTCCTCCGGAGGGCAGATTAGAACATGAGGATTGGAGATGCATGAAAC  **AS** use oligonucleotide from human pre-U8 |
| DNA template for production of human pre-U8 n.60G > C | **S**  TAATACGACTCACTATAGGGGATCGTCAGGTGGGATAATCCTTACCTGTTCCTCCTCCG  GAGGGCAGATTAGAACATGATCATTGGAGATGCATGAAAC  **AS** use oligonucleotide from human pre-U8 |
| DNA template for production of human pre-U8 n.60_61insT | **S** TAATACGACTCACTATAGGGGATCGTCAGGTGGGATAATCCTTACCTGTTCCTCCTCCGGAGGGCAGATTAGAACATGATGTATTGGAGATGCATG  **AS** use oligonucleotide from human pre-U8 |
| DNA template for production of human pre-U8 n.61A > G | PMID: 32359472 |
| DNA template for production of human pre-U8 n.61A > T | **S**  TAATACGACTCACTATAGGGGATCGTCAGGTGGGATAATCCTTACCTGTTCCTCCTCCGGAGGGCAGATTAGAACATGATGtTTGGAGATGCATG  **AS** use oligonucleotide from human pre-U8 |
| DNA template for production of human pre-U8 n.64G > A | **S**  TAATACGACTCACTATAGGGGATCGTCAGGTGGGATAATCCTTACCTGTTCCTCCTCCGGAGGGCAGATTAGAACATGATGATTAGAGATGCATGAAAC  **AS** use oligonucleotide from human pre-U8 |
| DNA template for production of human pre-U8 n.72A > G | **S**  use oligonucleotide from human pre-U8  **AS** ACAAATGTAAGTGATCGTCAGAAAGAATCAGACAGGAGCAATCAGGGTGTTGCAAGTCCTGATTACGCAGAGACGTTAATCACGTTTCACGCATCTCCAATCATC |
| DNA template for production of human pre-U8 n.73T > G | **S**  use oligonucleotide from human pre-U8  **AS** ACAAATGTAAGTGATCGTCAGAAAGAATCAGACAGGAGCAATCAGGGTGTTGCAAGTCCTGATTACGCAGAGACGTTAATCACGTTTCCTGCATCTCCAATCATC |
| DNA template for production of human pre-U8 n.74G > A | **S**  use oligonucleotide from human pre-U8  **AS** ACAAATGTAAGTGATCGTCAGAAAGAATCAGACAGGAGCAATCAGGGTGTTGCAAGTCCTGATTACGCAGAGACGTTAATCACGTTTTATGCATCTCCAATCATC |
| DNA template for production of human pre-U8 n.74G > T | **S**  use oligonucleotide from human pre-U8  **AS** ACAAATGTAAGTGATCGTCAGAAAGAATCAGACAGGAGCAATCAGGGTGTTGCAAGTCCTGATTACGCAGAGACGTTAATCACGTTTAATGCATCTCCAATCATC |
| DNA template for production of human pre-U8 n.75A > C | **S**  use oligonucleotide from human pre-U8  **AS** ACAAATGTAAGTGATCGTCAGAAAGAATCAGACAGGAGCAATCAGGGTGTTGCAAGTCCTGATTACGCAGAGACGTTAATCACGTTGCATGCATCTCCAATCATC |
| DNA template for production of human pre-U8 n.75A > G | **S**  use oligonucleotide from human pre-U8  **AS** ACAAATGTAAGTGATCGTCAGAAAGAATCAGACAGGAGCAATCAGGGTGTTGCAAGTCCTGATTACGCAGAGACGTTAATCACGTTCCATGCATCTCCAATCATC |
| DNA template for production of human pre-U8 n.81G > A | **S**  use oligonucleotide from human pre-U8  **AS** ACAAATGTAAGTGATCGTCAGAAAGAATCAGACAGGAGCAATCAGGGTGTTGCAAGTCCTGATTACGCAGAGACGTTAATTACGTTTCATGCATC |
| DNA template for production of human pre-U8 n.81G > C | **S**  use oligonucleotide from human pre-U8  **AS** ACAAATGTAAGTGATCGTCAGAAAGAATCAGACAGGAGCAATCAGGGTGTTGCAAGTCCTGATTACGCAGAGACGTTAATGACGTTTCATGCATC |
| DNA template for production of human pre-U8 n.82A > G | **S**  use oligonucleotide from human pre-U8  **AS** ACAAATGTAAGTGATCGTCAGAAAGAATCAGACAGGAGCAATCAGGGTGTTGCAAGTCCTGATTACGCAGAGACGTTAACCACGTTTCATGCATC |
| DNA template for production of human pre-U8 n.92C > T | **S**  use oligonucleotide from human pre-U8  **AS** ACAAATGTAAGTGATCGTCAGAAAGAATCAGACAGGAGCAATCAGGGTGTTGCAAGTCCTGATTACGCAAAGACGTTAATCACG |
| DNA template for production of human pre-U8 n.100T > G | **S**  use oligonucleotide from human pre-U8  **AS** ACAAATGTAAGTGATCGTCAGAAAGAATCAGACAGGAGCAATCAGGGTGTTGCAAGTCCTGCTTACGCAGAG |
| DNA template for production of human pre-U8 n.103G > A | **S**  use oligonucleotide from human pre-U8  **AS** ACAAATGTAAGTGATCGTCAGAAAGAATCAGACAGGAGCAATCAGGGTGTTGCAAGTCTTGATTACGCAGAG |
| DNA template for production of human pre-U8 n.104G > A | **S**  use oligonucleotide from human pre-U8  **AS** ACAAATGTAAGTGATCGTCAGAAAGAATCAGACAGGAGCAATCAGGGTGTTGCAAGTTCTGATTACGCAGAG |
| DNA template for production of human pre-U8 n.117C > G | **S**  use oligonucleotide from human pre-U8  **AS** ACAAATGTAAGTGATCGTCAGAAAGAATCAGACAGGAGCAATCACGGTGTTGCAAGTCCTG |
| DNA template for production of human pre-U8 n.118T > G | **S**  use oligonucleotide from human pre-U8  **AS** ACAAATGTAAGTGATCGTCAGAAAGAATCAGACAGGAGCAATCCGGGTGTTGCAAGTCCTG |
| DNA template for production of human pre-U8 n.126C > T | **S**  use oligonucleotide from human pre-U8  **AS** ACAAATGTAAGTGATCGTCAGAAAGAATCAGACAGAAGCAATCAGGGTGTTGCAAG |
| DNA template for production of human pre-U8 n.127C > G | **S**  use oligonucleotide from human pre-U8  **AS** ACAAATGTAAGTGATCGTCAGAAAGAATCAGACACGAGCAATCAGGGTGTTGCAAG |
| DNA template for production of human pre-U8 n.130T > C | **S**  use oligonucleotide from human pre-U8  **AS** ACAAATGTAAGTGATCGTCAGAAAGAATCAGGCAGGAGCAATCAG |
| DNA template for production of human pre-U8 n.131C > A | **S**  use oligonucleotide from human pre-U8  **AS** ACAAATGTAAGTGATCGTCAGAAAGAATCATACAGGAGCAATCAG |
| DNA template for production of human pre-U8 n.131C > G | **S**  use oligonucleotide from human pre-U8  **AS** ACAAATGTAAGTGATCGTCAGAAAGAATCACACAGGAGCAATCAG |
| DNA template for production of human pre-U8 n.131C > T | **S**  use oligonucleotide from human pre-U8  **AS** ACAAATGTAAGTGATCGTCAGAAAGAATCAAACAGGAGCAATCAG |
| DNA template for production of human pre-U8 n.*1C > T | PMID: 32359472 |
| DNA template for production of human pre-U8 n.*5C > G | PMID: 32359472 |
| DNA template for production of human pre-U8 n.*9C > T | PMID: 32359472 |
| DNA template for production of human pre-U8 n.*10G > A | **S**  use oligonucleotide from human pre-U8  **AS** ACAAATGTAAGTGATtGTCAgAAAGAATCAGACAGGAG |
| DNA template for production of human pre-U8 n.*10G > T | PMID: 32359472 |
| DNA template for production of human pre-U8 n.21T > G | **S**  TAATACGACTCACTATAGGGGATCGTCAGGTGGGATAATCCGTACCTGTTCCTCCTC  **AS** use oligonucleotide from human pre-U8 |
| DNA template for production of human pre-U8 n.108T > G | **S**  use oligonucleotide from human pre-U8  **AS** ACAAATGTAAGTGATCGTCAGAAAGAATCAGACAGGAGCAATCAGGGTGTTGCCAGTCCTGATTACGCAGAG |
| DNA template for production of human pre-U8 n.53A > G | **S** TAATACGACTCACTATAGGGGATCGTCAGGTGGGATAATCCTTACCTGTTCCTCCTCCGGAGGGCAGATTAGAGCATGATGATTGGAGATGCATG  **AS** use oligonucleotide from human pre-U8 |
| DNA template for production of human pre-U8 n.105A > T | **S**  use oligonucleotide from human pre-U8  **AS** ACAAATGTAAGTGATCGTCAGAAAGAATCAGACAGGAGCAATCAGGGTGTTGCAAGACCTGATTACGCAGAG |
| DNA template for production of human pre-U8 n.112A > G | **S**  use oligonucleotide from human pre-U8  **AS** ACAAATGTAAGTGATCGTCAGAAAGAATCAGACAGGAGCAATCAGGGTGCTGCAAGTCCTGATTACGCAGAG |
| DNA template for production of human pre-U8 n.110C > T | **S**  use oligonucleotide from human pre-U8  **AS** ACAAATGTAAGTGATCGTCAGAAAGAATCAGACAGGAGCAATCAGGGTGTTACAAGTCCTGATTACGCAGAG |
| DNA template for production of human pre-U8 n.113C > T | **S**  use oligonucleotide from human pre-U8  **AS** ACAAATGTAAGTGATCGTCAGAAAGAATCAGACAGGAGCAATCAGGGTATTGCAAGTCCTGATTACGCAGAG |
| DNA template for production of human pre-U8 n.110C > T; n.113C > T | **S**  use oligonucleotide from human pre-U8  **AS** ACAAATGTAAGTGATCGTCAGAAAGAATCAGACAGGAGCAATCAGGGTATTACAAGTCCTGATTACGCAGAG |
| DNA template for production of zebrafish pre-U8.3 n.35G > C | **S**  TAATACGACTCACTATAGGGGACATCAGTGAGGTACATCCTTACCTGTTACCTTACCAATAAGGTTACAGATCATG  **AS** use oligonucleotide from zebrafish pre-U8.3 |
| DNA template for production of gRNA to generate 47bp hypomorphic mutation | PMID: 32359472 |
