## Supplementary figures for "Human Mendelian disease and *in vivo* mutagenesis screening define the molecular architecture of the U8 snoRNA"

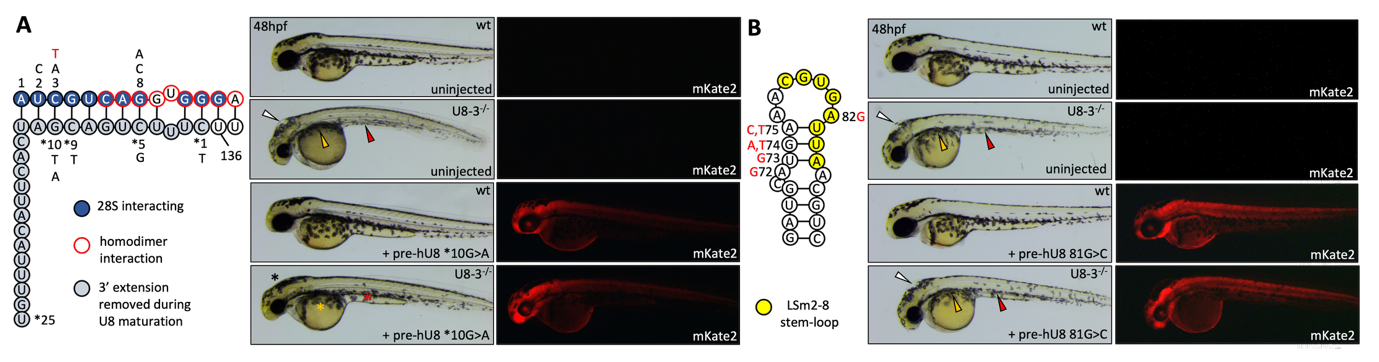

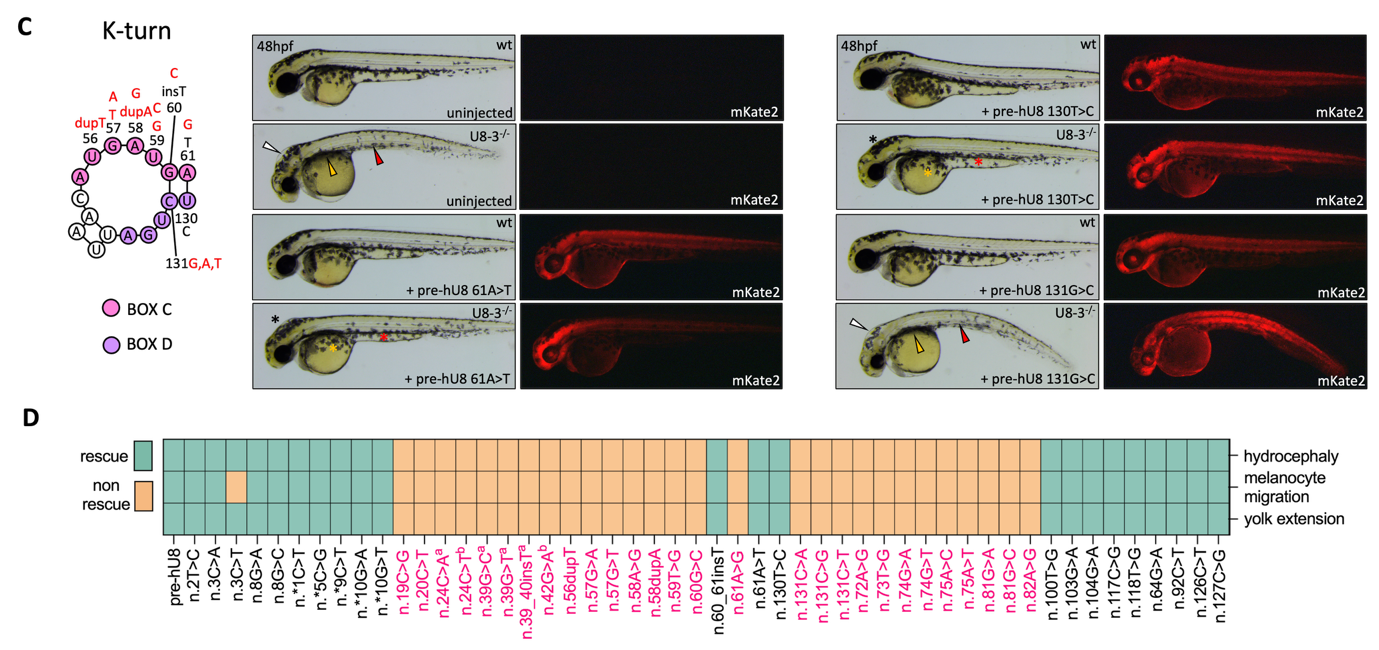

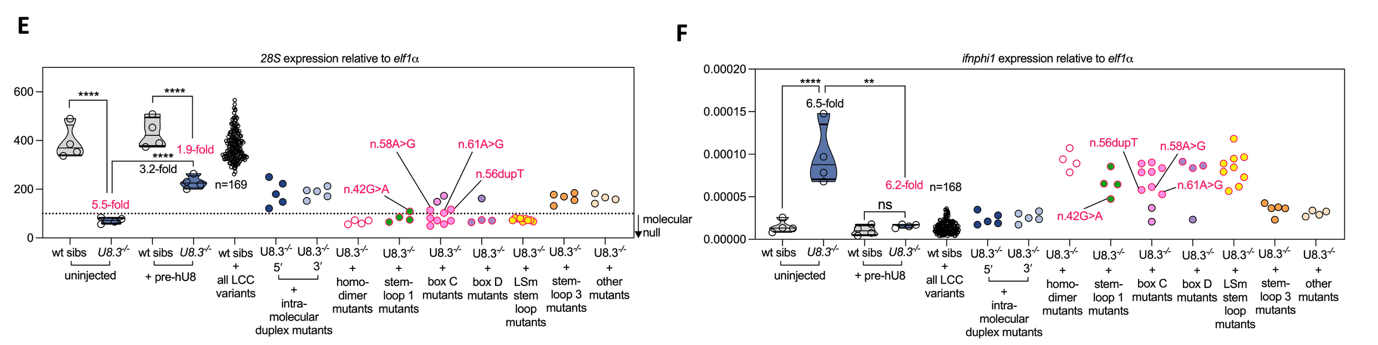

**Figure S1** **A zebrafish bioassay characterises LCC-causative U8 mutations into hypomorphic and null alleles based upon morphological rescue of the U8.3 mutant. A**.) Schematic depicting the intramolecular duplex that forms between the 5′ end and 3′ extension of precursor U8, and the LCC-causative mutations that occur in this region. Representative brightfield and fluorescent images of the indicated genotypes, exogenous snoRNA and fluorescent protein taken at 48hpf. **B**.) Schematic depicting stem-loop structure containing the LSm-binding octamer, and the LCC-causative mutations that occur in this region. Representative brightfield and fluorescent images of the indicated genotypes, exogenous snoRNA and fluorescent protein taken at 48hpf. **C**.) Schematic depicting the K-turn that forms between the Box C and Box D motifs, and the LCC-causative mutations that occur in this region. Representative brightfield and fluorescent images of the indicated genotypes, exogenous snoRNAs and fluorescent protein taken at 48hpf. **D**.) Green (positively) or orange (negatively) indicate whether the hydrocephaly, melanocyte migration or yolk extension defects seen in the U8.3^-/-^ mutant embryo are rescued by the indicated U8 snoRNA. **E**.) qRT-PCR of *28S* for the indicated snoRNAs at 48hpf in the noted genotypes. Variants below the dashed line represent null alleles. U8 mutants that rescue the morphology of the U8.3^-/-^ mutant more effectively increase 28S levels. n.42G > A, n.56dupT, n.58A > G and n.61A > G mutants confer residual function in 28S biogenesis that is insufficient to rescue the morphology of U8.3^-/-^ embryos at 48hpf. **F**.) qRT-PCR of *ifnphi1* for the indicated snoRNAs at 48hpf in the given genotypes. Variants that rescue the morphology of the U8.3^-/-^ mutant more effectively suppress *ifnphi1* induction.

White, red and orange arrowheads denote, respectively, hydrocephaly, aberrant yolk extension and impaired migration of melanocytes over the yolk. Black, red and orange asterisks denote, respectively, rescued hindbrain, yolk extension and melanocyte migration. Significance determined by unpaired t-test. Each biological replicate is indicated by a circle, except for LCC-causative U8 mutant snoRNAs, which are mean values of 3-6 biological replicates. Black and red fold change indicates increase and decrease respectively. All *hU8* variants were microinjected as precursors. For panels E-F, *hU8* mutants that rescue the morphology of the U8.3^-/-^ embryo are outlined in black, and *hU8* mutants that fail to rescue U8.3^-/-^ embryo morphology outlined in red.

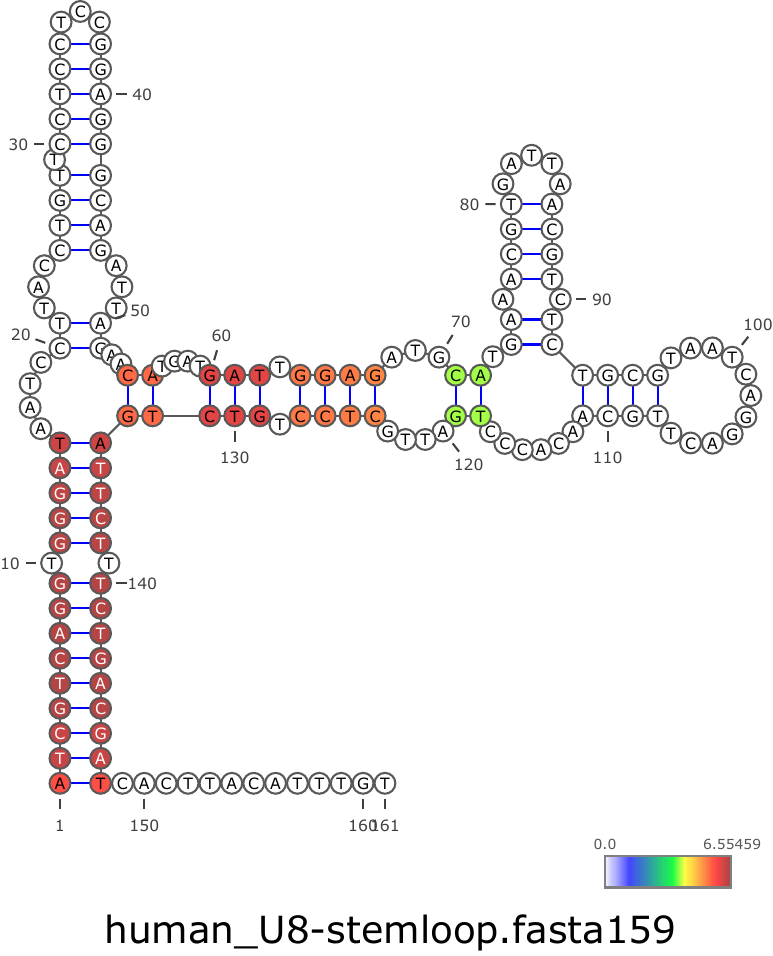

**Figure S2** **Analysis of the PARIS2 dataset confirms that an intramolecular duplex forms between the 5′ end of U8 and its 3′ extension in human cells.** Schematic illustration of pre-U8 secondary structure obtained from the PARIS2 dataset with confidence predictions for the stem-loop structures shown.

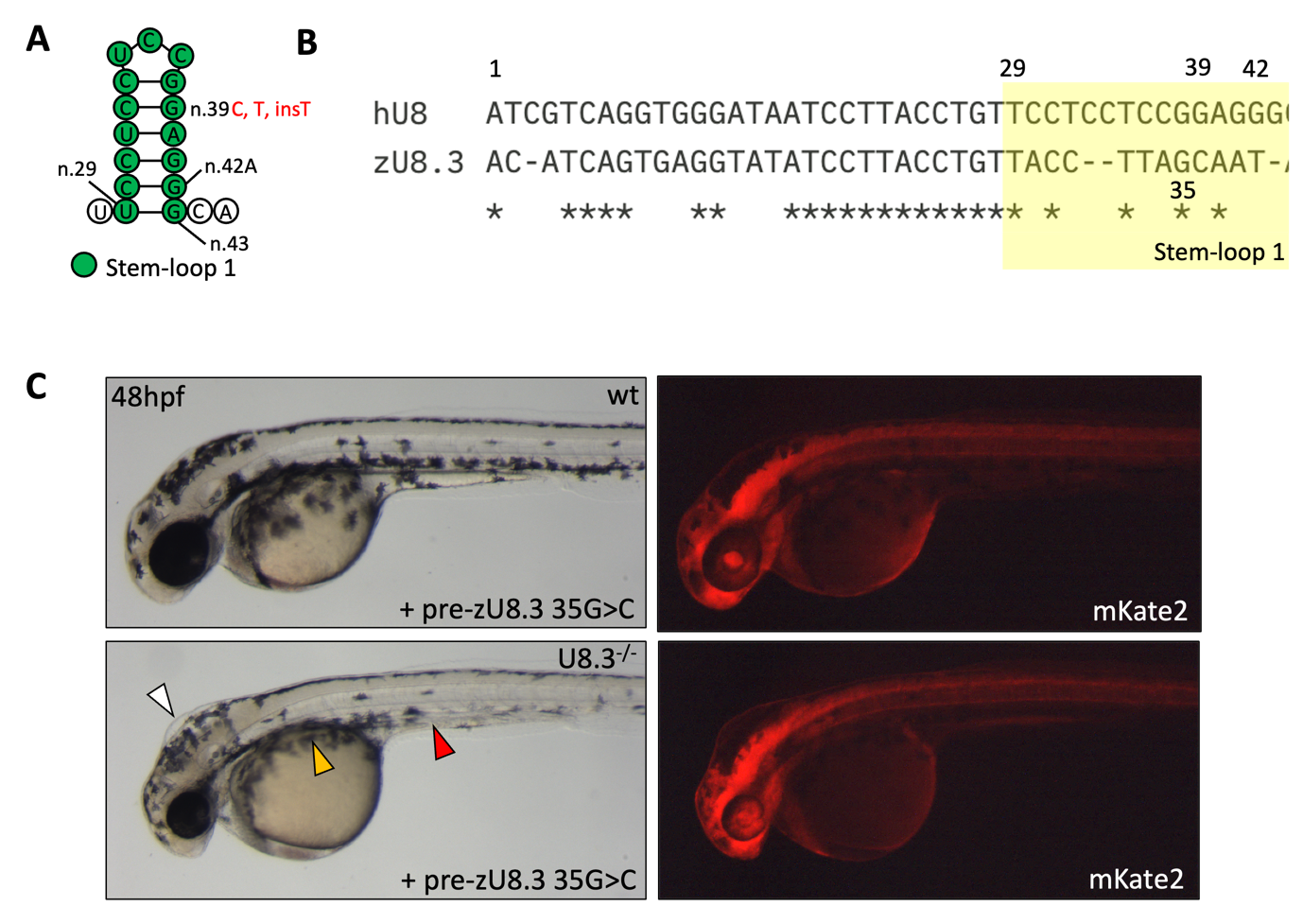

**Figure S3** **Stem-loop 1 of mature U8 is poorly conserved in zebrafish. A**.) Schematic depicting stem-loop 1 of mature human U8, and the LCC-causative mutations that occur in this region. **B**.) Alignment of U8 stem-loop 1 sequence (yellow box) demonstrates poor conservation between human and zebrafish U8-3. **C**.) Representative brightfield and fluorescent images of the indicated genotypes, exogenous snoRNA and fluorescent protein taken at 48hpf. White, red and orange arrowheads denote, respectively, hydrocephaly, aberrant yolk extension and impaired migration of melanocytes over the yolk. Mutation of a single G nucleotide within stem-loop 1 is sufficient to inactive zebrafish U8.3.

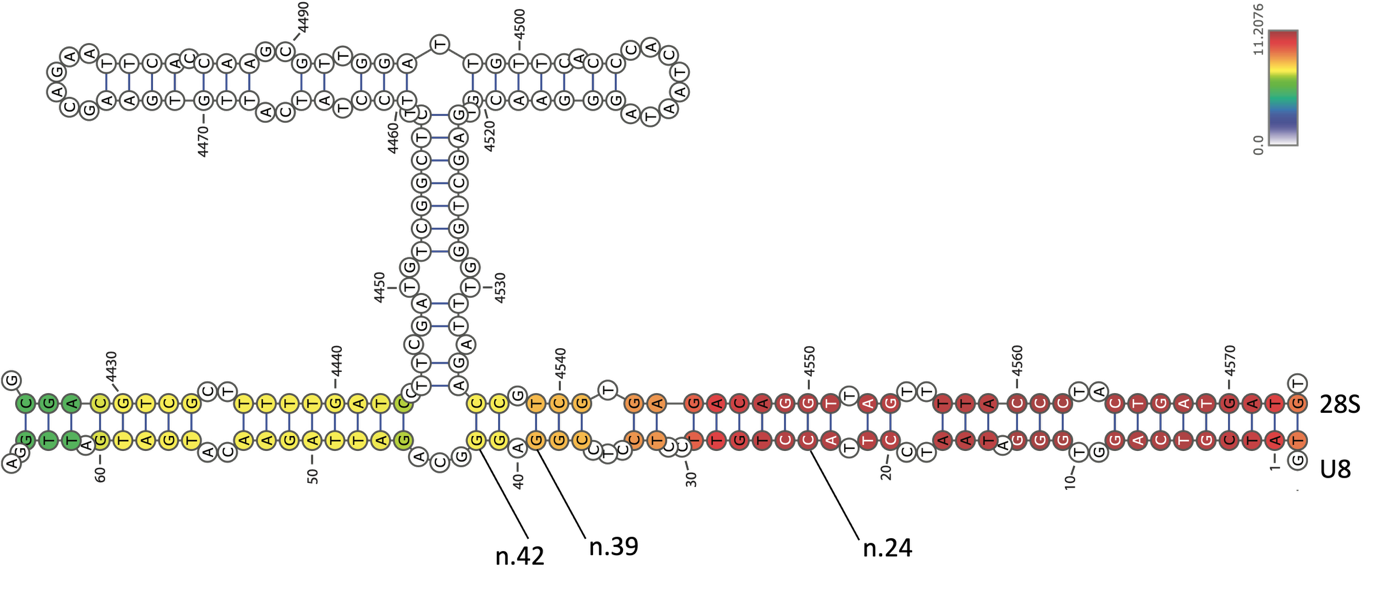

**Figure S4** **28S-U8 duplex formation obtained from the PARIS2 dataset.** Schematic illustration of the interaction between 28S and human U8 with confidence predictions shown. n.19 of U8 is not predicted to interact with 28S, whereas n.24 is predicted to interact with 28S with high confidence. Lower confidence interactions of n.39 and n.42 of U8 with 28S are predicted.

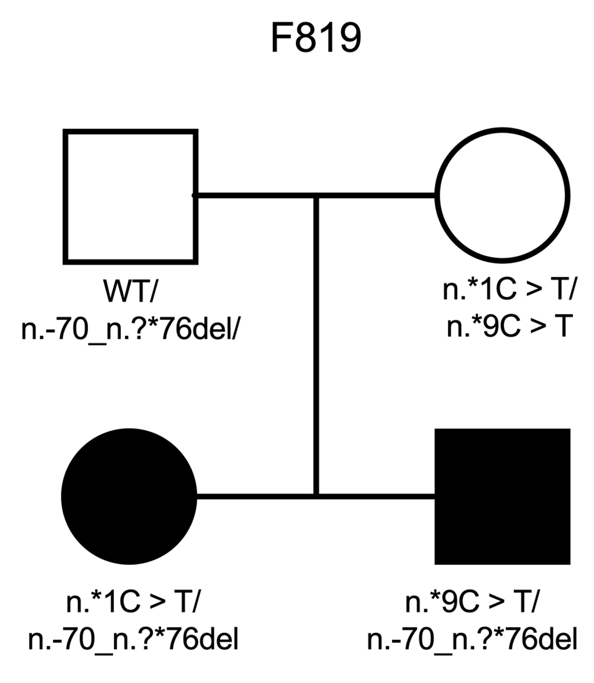

**Figure S5 Family pedigree of F819**. Note that while both affected children inherited the same paternal whole gene deletion, they each inherited a different hypomorphic mutation affecting processing of pre-U8 from their asymptomatic mother.

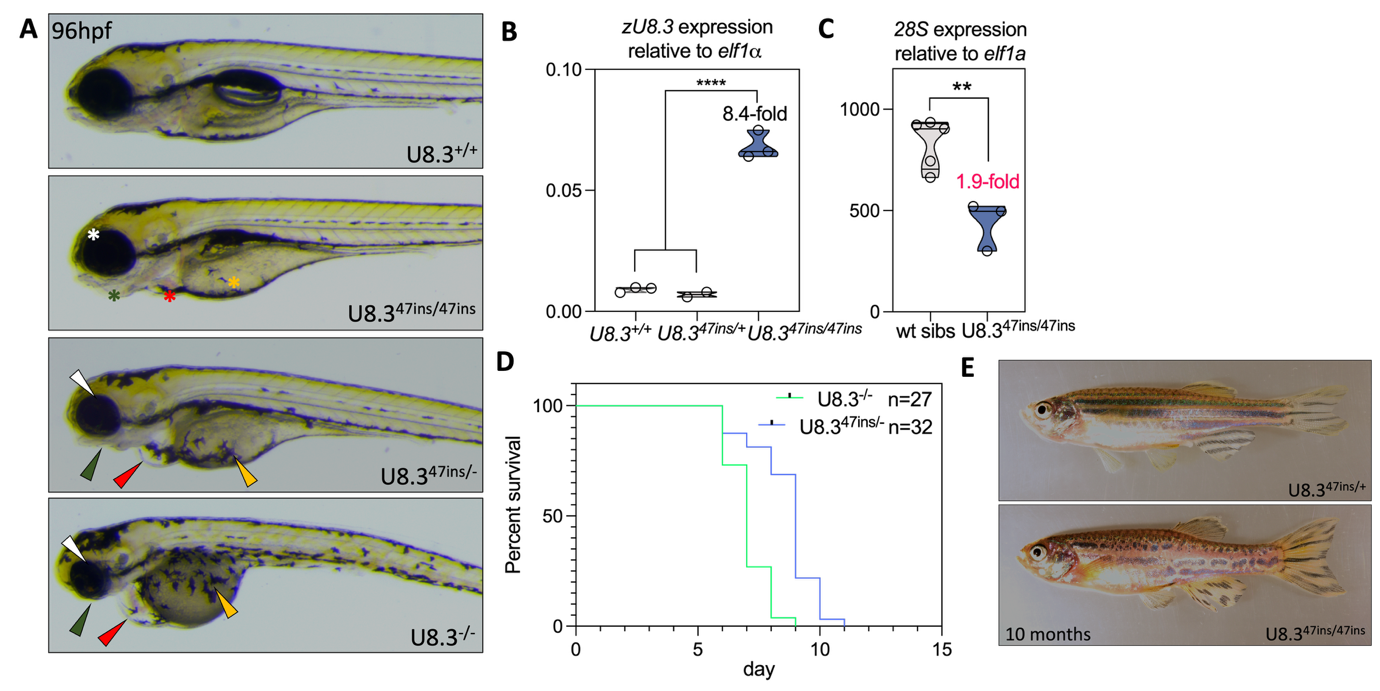

**Figure S6** **Zebrafish homozygous for a hypomorphic U8.3 allele are adult viable, demonstrating that U8.3 is subject to a gene dosage effect.** **A**.) Representative brightfield images of the indicated genotypes. U8.3^47ins/-^ and U8.3^-/-^ larvae 96hpf exhibit reduced eye size (white arrowhead), reduced jaw development (green arrowhead), cardiac oedema (red arrowhead) and delayed melanocyte migration over the yolk (orange arrowhead) when compared to U8.3^+/+^ larvae. These morphological abnormalities are rescued in U8.3^47ins/47ins^ larvae (white, green, red, and orange asterisks respectively). U8.3^47ins/47ins^ larvae are developmentally delayed, and inflate their swim bladder at 120hpf, rather than at 96hpf in U8.3^+/+^ larvae. **B**.) qRT-PCR of z*U8.3* at 96hpf in the indicated genotypes. **C**.) qRT-PCR of *28S* at 96hpf in the indicated genotypes. **D**.) Kaplan-Meier plot indicating that the U8.3^47ins/-^ and U8.3^-/-^ genotypes are lethal. **E**.) Representative images showing U8.3^47ins/47ins^ animals are adult viable, fertile and display abnormal melanocyte pigmentation.

Significance determined by unpaired t-test. Each biological replicate is indicated by a circle. Red fold change indicates decrease, black fold change indicates increase. The 47bp hypomorphic allele is denoted by 47ins.

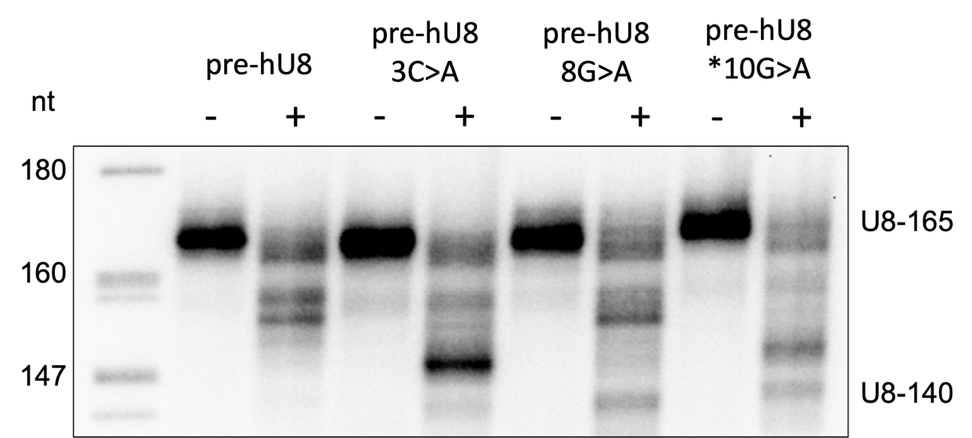

**Figure S7 LCC-associated U8 mutations increase the rate of processing of precursor U8.** Processing of 5′ end-radiolabelled *in vitro* transcribed precursor U8 wildtype/mutant snoRNA U8-165 was assessed in (+) or without (-) HeLa nuclear extracts. At 30 min, 3C>A, 8G>A and *10G>A mutant snoRNA exhibit an enhanced rate of processing when compared to wildtype, as demonstrated by the presence of mature U8 (U8-140).

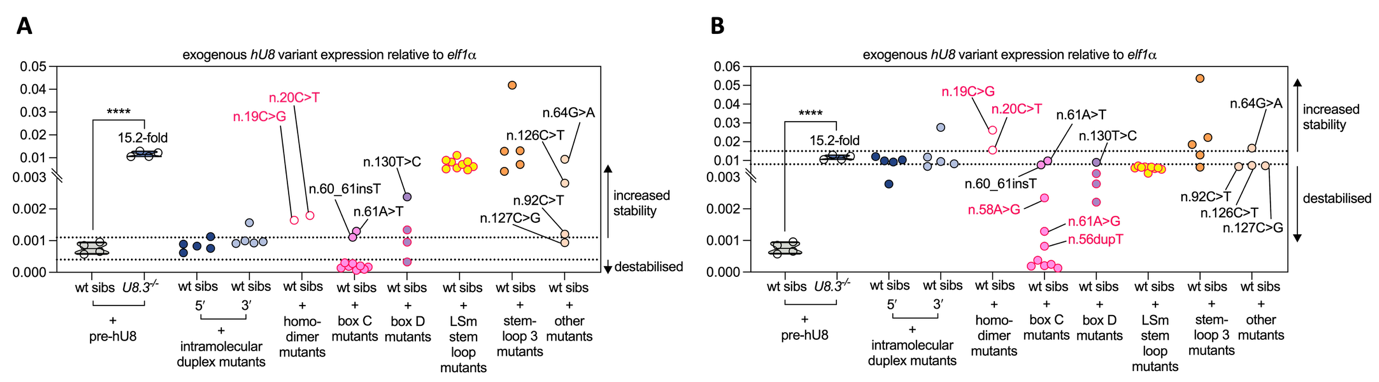

**Figure S8 The effect of** **LCC-causative mutations on U8 stability clusters according to U8 structural and protein interacting domains. A**.) qRT-PCR of the indicated snoRNAs at 48hpf in the given genotypes. The dashed lines represent the range of expected stability of wildtype *hU8* in wildtype sibling embryos. **B**.) qRT-PCR of the indicated snoRNAs at 48hpf in the given genotypes. The dashed lines represent the range of expected stability of wildtype *hU8* in U8.3^-/-^ embryos.

Significance determined by unpaired t-test. Each biological replicate is indicated by a circle, except for LCC-causative U8 mutant snoRNAs which are mean values of 3-6 biological replicates. Black fold change indicates increase. All *hU8* variants were microinjected as precursors. *hU8* mutants that rescue the morphology of the U8.3^-/-^ embryo are outlined in black, and *hU8* mutants that fail to rescue U8.3^-/-^ embryo morphology outlined in red. Box C mutations with residual function in 28S biogenesis (n.56dupT, n.58A > G and n.61A > G) exhibit elevated stability in U8.3^-/-^ embryos compared to box C mutants with null function.

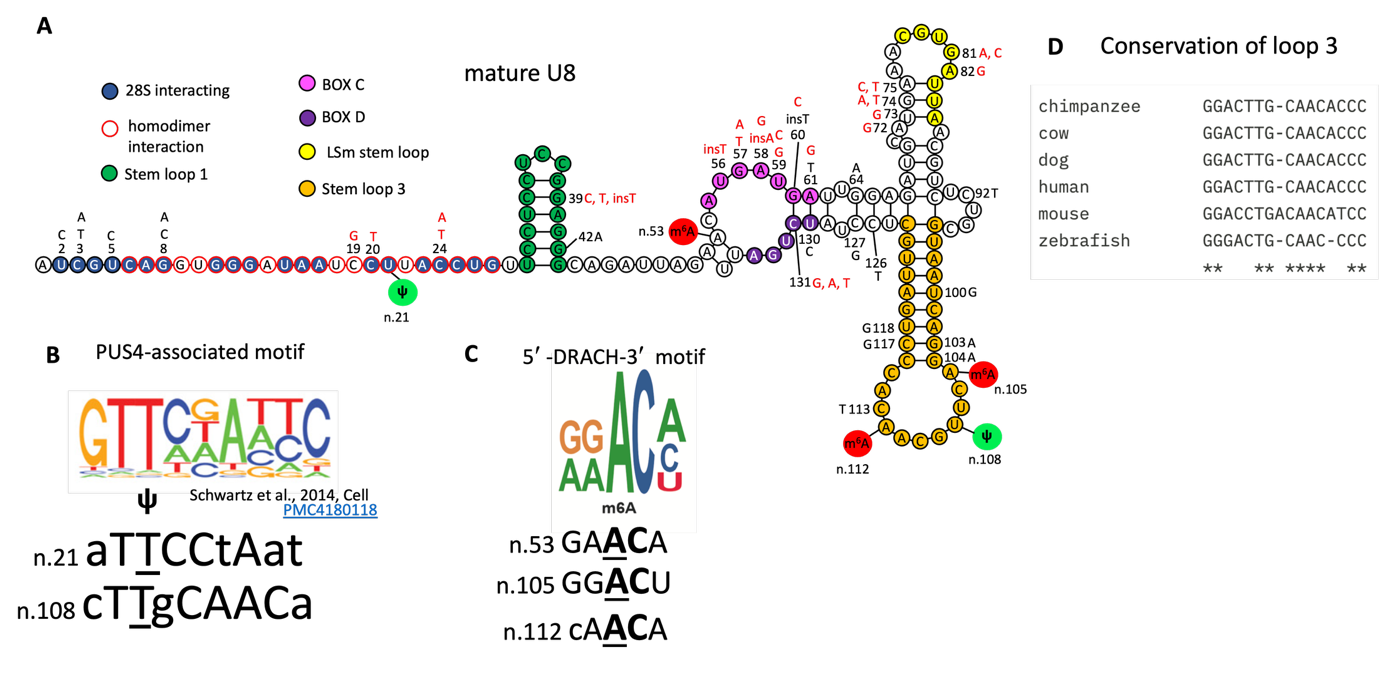

**Figure S9** **Human U8 is modified at five nucleotides within PUS4 and DRACH motifs. A**.) Schematic depicting the secondary structure of mature U8 snoRNA. Mutations that rescue or fail to rescue U8.3^-/-^ mutant morphology are coloured black or red respectively. The position of pseudouridine modifications (Ψ) are shown in green, and N-6-methylation modifications (m^6^A) in red. **B**.) Pseudouridine modifications occur in PUS4-associated motifs. **C**.) n.53A and n.105A m^6^A modifications in U8 conform to a DRACH motif, whereas n.112A is divergent in the most 5′ nucleotide. **D**.) Multiple species alignment of the loop sequence of stem-loop 3 of U8.

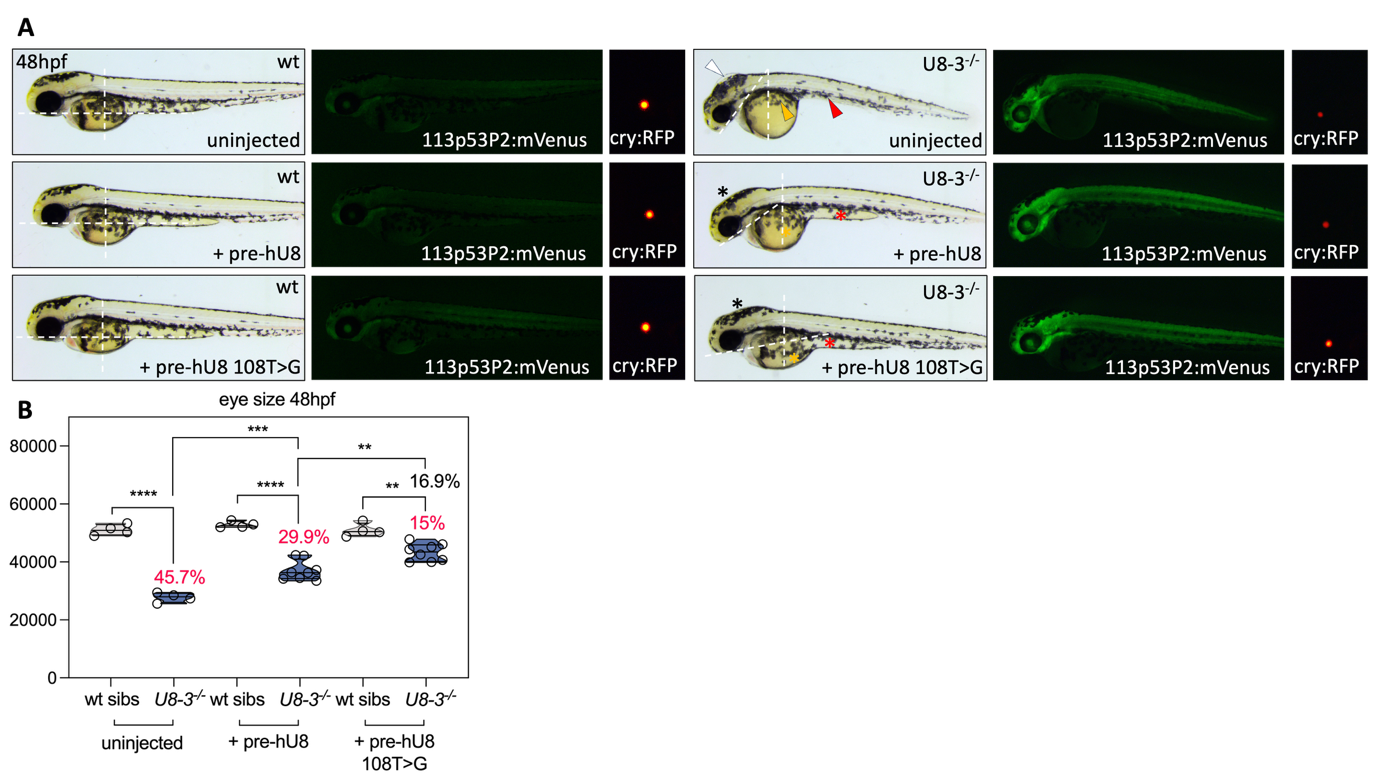

**Figure S10** **Mutation of n.108 enhances the function of U8 A**.) Representative brightfield and fluorescent images of the indicated genotypes, exogenous snoRNAs and fluorescent protein is given at 48hpf. Tp53 activity is reported under control of the 113p53P2 promoter that drives expression of mVenus, and was used to identify rescued mutants. The integrated 113p53P2:mVenus transgene is reported by γ-crystallin:mCherry which drives lens specific expression of mCherry from approximately 36hpf. The dashed white lines indicate the angle of the head relative to the yolk, with developmentally older embryos closer to 90°C at 48hpf. **B**.) The left eye area is quantified for the indicated genotypes microinjected with the given exogenous U8 snoRNAs.

White, red and orange arrowheads denote, respectively, hydrocephaly, aberrant yolk extension and impaired migration of melanocytes over the yolk. Black, red and orange asterisks denote, respectively, rescued hindbrain, yolk extension and melanocyte migration. pre - precursor. Significance determined by unpaired t-test. Each biological replicate is indicated by a circle. Red percentage indicates the amount the mutant eye is decreased from wildtype; black percentage indicates the amount the U8.3 mutant eye injected with 108T>G is increased from that of the U8.3 mutant eye injected with pre-U8.

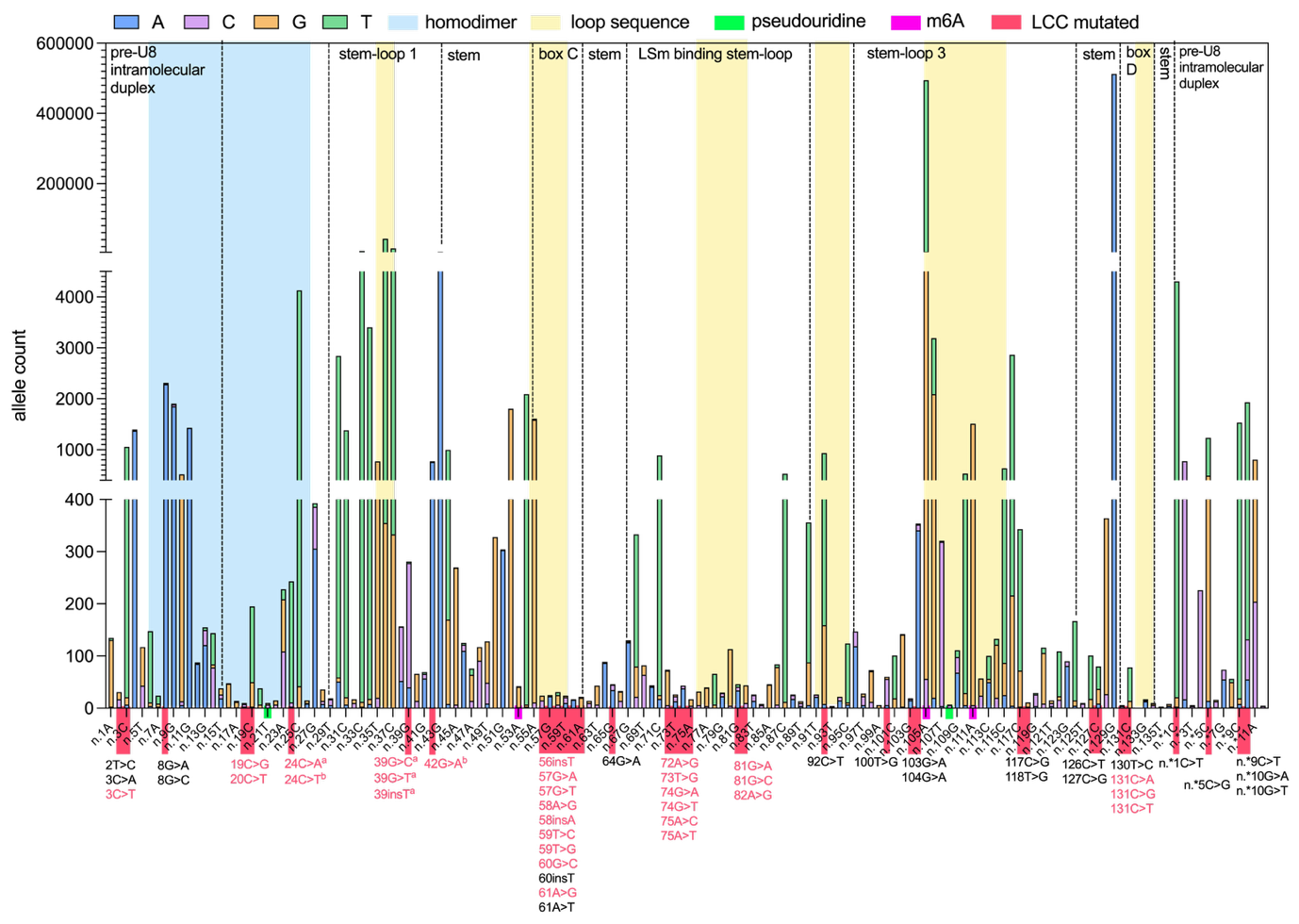

**Figure S11** **Human allelic variation in U8 is highest in regions known to mediate interaction with other RNA molecules, and is constrained in regions that interact with protein or stem structures.** Total allele counts for each nucleotide of precursor U8 is shown from the Genome Aggregation Database (gnomAD) v.4.1.0. The position of LCC-causative mutations is highlighted by red boxes, and the functionality of these mutations is recorded in red (null) or black (hypomorphic) according to human genetic and zebrafish bioassay data.

^a^denotes U8 mutants that are detected in the homozygous state in patients with LCC (n.39G > C, n.39G >T), or in the gnomAD v4.1.0 database (n.39_40insT), or co-occur with known null alleles in patients with LCC (n.24C > A co-occurs with n.82A >G), indicating hypomorphic function since homozygous null alleles are incompatible with life.

^b^denotes U8 mutants whose functionality is uncertain due to evidence that the affected nucleotides localise to regions of U8 that are not conserved at the sequence level in zebrafish. The effect on U8 stability of mutations denoted ^a^ or ^b^ is not included due to this lack of conservation.

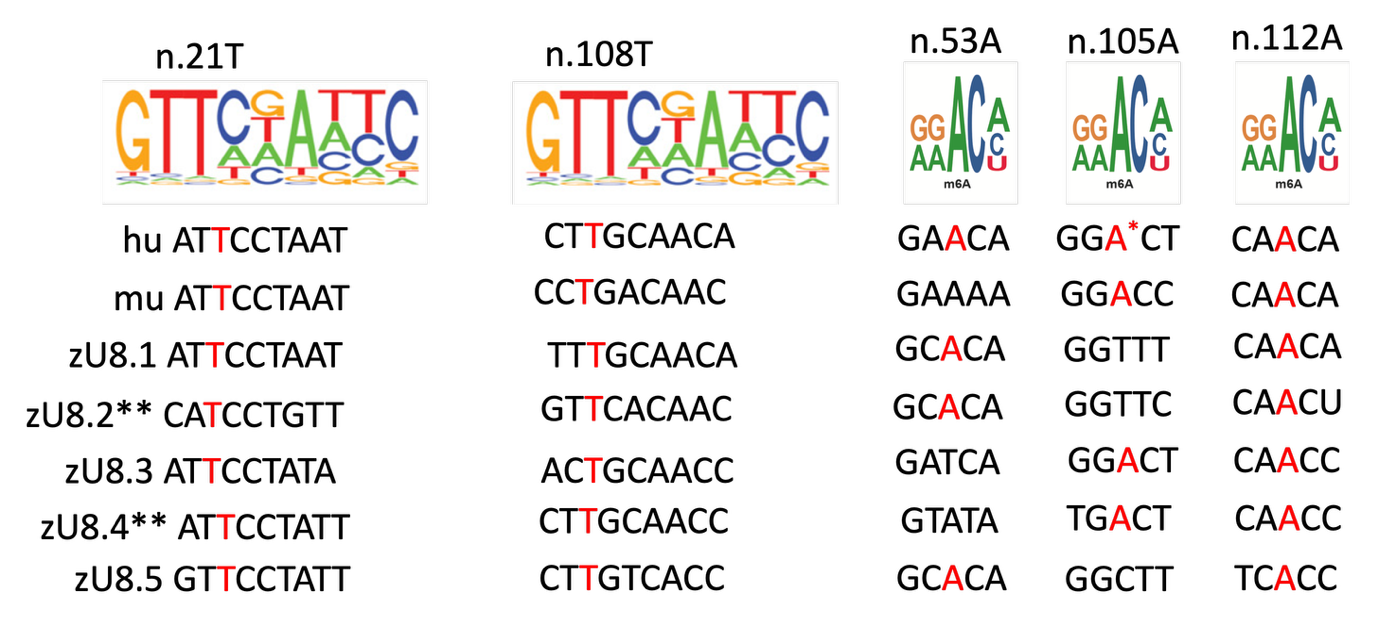

**Figure S12** **Conservation of U8 modified sites across human, mouse and zebrafish.** Conservation of U8 modification sites between human, mouse and zebrafish. Putatively modified nucleotides are highlighted in red, except where modifications are not predicted.

*Predominant variant is n.105T in human.

**zU8.2 and zU8.4 exhibit significant sequence variation in the LSm-binding and D box motifs respectively, and are likely to be non-functional pseudogenes.
