## Supplementary tables for "Human Mendelian disease and *in vivo* mutagenesis screening define the molecular architecture of the U8 snoRNA"

| **U8 variant /**  **Genomic coordinate** | **gnomAD**  **v4.1.0 classification** | **Functional classification**  **(this study)** | **Reasoning** |
| --- | --- | --- | --- |
| n.2T > C /  Chr17:8173587 A > G | Pathogenic | Hypomorph | Rescues U8 zebrafish mutant, alters processing of precursor U8 |
| n.3C > A /  Chr17:8173586 G > T | Pathogenic | Hypomorph | Rescues U8 zebrafish mutant, alters processing of precursor U8 |
| n.3C > T /  Chr17:8173586 G > A | Conflicting interpretation | Hypomorph | 2 homozygous individuals gnomAD, rescues U8 zebrafish mutant, alters processing of precursor U8 |
| n.8G > A /  Chr17:8173581 C > T | Benign/Likely benign | Hypomorph | Rescues U8 zebrafish mutant, 4 homozygous individuals gnomAD, alters processing of precursor U8 |
| n.8G > C /  Chr17:8173581 C > G | Pathogenic/Likely pathogenic | Hypomorph | Rescues U8 zebrafish mutant, homozygous in LCC patient, alters processing of precursor U8 |
| n.19C > G /  Chr17:8173570 G > C | Uncertain significance | Amorph | Fails to rescue U8 zebrafish mutant |
| n.20C > T /  Chr17:8173569 G > A | Uncertain significance | Amorph | Fails to rescue U8 zebrafish mutant |
| n.24C > A^a^ /  Chr17:8173565 G > T | Uncertain significance | Hypomorph | Observed in combination with known null allele (n.82A > G). Two null mutations are incompatible with life, indicating n.24C > A must be functional |
| n.24C > T^b^ /  Chr17:8173565 G > A | Conflicting interpretation | Unknown | Fails to rescue U8 zebrafish mutant, co-occurs with hypomorphic n.*10G > T and n.3C >T in two different patients. Null or hypomorphic function is possible. |
| n.39G > C^a^ /  Chr17:8173550 C > G | Pathogenic | Hypomorph | Homozygous in LCC patient |
| n.39G > T^a^ /  Chr17:8173550 C > A | Pathogenic | Hypomorph | Homozygous in LCC patient |
| n.39_40insT^a^ /  Chr17: 8173549_8173550insA | Unclassified | Hypomorph | 1 individual homozygous in gnomAD |
| n.42G > A^b^ /  Chr17:8173547 C > T | Conflicting interpretation | Unknown | Fails to rescue the U8 zebrafish mutant, but residual function at molecular level. Localises to stem-loop 1, which is poorly conserved in zebrafish. Co-occurs with hypomorphic n.3C > A. Null or hypomorphic function is possible |
| n.56dupT /  Chr17:8173532 dup | Pathogenic | Amorph | Fails to rescue U8 zebrafish mutant |
| n.57G > A /  Chr17:8173532 C > T | Uncertain significance | Amorph | Fails to rescue U8 zebrafish mutant |
| n.57G > T /  Chr17:8173532 C > A | Pathogenic | Amorph | Fails to rescue U8 zebrafish mutant |
| n.58A > G /  Chr17:8173531 T > C | Uncertain significance | Amorph | Fails to rescue U8 zebrafish mutant |
| n.58dupA /  Chr17:8173530 dup | Not present in database | Amorph | Fails to rescue U8 zebrafish mutant |
| n.59T > G /  Chr17:8173530 A > C | Pathogenic | Amorph | Fails to rescue U8 zebrafish mutant |
| n.60G > C /  Chr17:8173529 C > G | Not present in database | Amorph | Fails to rescue U8 zebrafish mutant |
| n.60_61insT /  Chr17:8173528 _8173529 insA | Pathogenic | Hypomorph | Rescues U8 zebrafish mutant |
| n.61A > G /  Chr17:8173528 T > C | Uncertain significance | Amorph | Fails to rescue U8 zebrafish mutant |
| n.61A > T /  Chr17:8173528 T > A | Unclassified | Hypomorph | Rescues U8 zebrafish mutant |
| n.64G > A /  Chr17:8173525 C > T | Uncertain significance | Hypomorph | Rescues U8 zebrafish mutant |
| n.72A > G /  Chr17:8173517 T > C | Pathogenic/Likely pathogenic | Amorph | Fails to rescue U8 zebrafish mutant |
| n.73T > G /  Chr17:8173516 A > C | Pathogenic | Amorph | Fails to rescue U8 zebrafish mutant |
| n.74G > A /  Chr17:8173515 C > T | Uncertain significance | Amorph | Fails to rescue U8 zebrafish mutant |
| n.74G > T /  Chr17:8173515 C > A | Not present in database | Amorph | Fails to rescue U8 zebrafish mutant |
| n.75A > C /  Chr17:8173514 T > G | Pathogenic | Amorph | Fails to rescue U8 zebrafish mutant |
| n.75A > T /  Chr17:8173514 T > A | Not present in database | Amorph | Fails to rescue U8 zebrafish mutant |
| n.81G > A /  Chr17:8173508 C > T | Likely pathogenic | Amorph | Fails to rescue U8 zebrafish mutant |
| n.81G > C /  Chr17:8173508 C > G | Pathogenic | Amorph | Fails to rescue U8 zebrafish mutant |
| n.82A > G /  Chr17:8173507 T > C | Pathogenic | Amorph | Fails to rescue U8 zebrafish mutant |
| n.92C > T /  Chr17:8173497 G > A | Likely benign | Hypomorph | Rescues U8 zebrafish mutant, observed in combination with null 72A>G allele |
| n.100 T > G /  Chr17:8173489 A > C | Pathogenic | Hypomorph | Rescues U8 zebrafish mutant |
| n.103G > A /  Chr17:8173486 C > T | Uncertain significance | Hypomorph | Rescues U8 zebrafish mutant |
| n.104G > A /  Chr17:8173485 C > T | Conflicting interpretation | Hypomorph | Rescues U8 zebrafish mutant |
| n.117C > G /  Chr17:8173472 G > C | Unclassified | Hypomorph | Rescues U8 zebrafish mutant |
| n.118T > G /  Chr17:8173471 A > C | Unclassified | Hypomorph | Rescues U8 zebrafish mutant |
| n.126C > T /  Chr17:8173463 G > A | Uncertain significance | Hypomorph | Rescues U8 zebrafish mutant |
| n.127C > G /  Chr17:8173462 G > C | Pathogenic | Hypomorph | Rescues U8 zebrafish mutant |
| n.130T > C /  Chr17:8173459 A > G | Pathogenic | Hypomorph | Rescues U8 zebrafish mutant |
| n.131C > A /  Chr17:8173458 G > T | Pathogenic | Amorph | Fails to rescue U8 zebrafish mutant |
| n.131C > G /  Chr17:8173458 G > C | Conflicting interpretation | Amorph | Fails to rescue U8 zebrafish mutant |
| n. 131C> T /  Chr17:8173458 G > A | Unclassified | Amorph | Fails to rescue U8 zebrafish mutant |
| n.*1C > T /  Chr17:8173452 G > A | Conflicting interpretation | Hypomorph | Rescues U8 zebrafish mutant, 5 homozygous individuals gnomAD, alters processing of precursor U8 |
| n.*5C > G /  Chr17:8173448 G > C | Pathogenic | Hypomorph | Rescues U8 zebrafish mutant, homozygous in LCC patients, alters processing of precursor U8 |
| n.*9C > T /  Chr17:8173444 G > A | Pathogenic/Likely pathogenic | Hypomorph | Rescues U8 zebrafish mutant, 2 homozygous individuals gnomAD, alters processing of precursor U8 |
| n.*10G > A /  Chr17:8173443 C > T | Likely pathogenic | Hypomorph | Rescues U8 zebrafish mutant, alters processing of precursor U8 |
| n.*10G > T /  Chr17:8173443 C > A | Benign | Hypomorph | Rescues U8 zebrafish mutant, 4 homozygous individuals gnomAD,  alters processing of precursor U8 |

**Table S1** **Human Mendelian genetics and zebrafish bioassay data classify LCC-associated U8 mutations into hypomorphic and null** **alleles.**

In column 1, hypomorphic mutations are coloured black and null mutations in red. Evidence for allelic classification and pathogenicity is summarised in column 4.

^a^denotes U8 mutants that are detected in homozygous form in LCC patients (n.39G > C, n.39G >T), or the gnomad v4.1.0 database (n.39_40insT), or co-occur with known null alleles in LCC patients (n.24C > A co-occurs with n.82A >G), indicating hypomorphic function despite an inability to rescue the U8.3^-/-^ mutant phenotype.

^b^denotes U8 mutants whose functionality is uncertain due to evidence that the affected nucleotide localises to regions of human U8 that are not functionally conserved in the zebrafish bioassay.

| **Crow cohort** |
| --- |

| **Patient** | **Age at onset** | | **Genotype** | | | | **Compound heterozygous hypomorphic alleles** |
| --- | --- | --- | --- | --- | --- | --- | --- |
| F172 | 8 W | | n.56dupT/n.*10G > T | | | |  |
| F278 | 1 Y | | n.8G > C/n.75A > G | | | |  |
| F281 | 4 Y | | n.3C > T/n.81G > C | | | |  |
| F285 | 12 Y | | n.57G > A/n.*5C > G | | | |  |
| F309 | 2 Y | | n.3C > T/n.131C > G | | | |  |
| F330* | 3 Y | | *n.25C > T^a^*/n.82A > G | | | |  |
| F331.1 | 2 Y | | n.72A > G/n.*10G > T | | | |  |
| F331.2 | 5 Mo | | n.72A > G/n.*10G > T | | | |  |
| F334 | 6 Y | | n.82A > G/n.*10G > T | | | |  |
| F337 | 18 Mo | | n.2T > C/n.58dupA | | | |  |
| F343 | 9 Y | | *n.-7_22dup* /n.*5C > G | | | |  |
| F344 | 7 W | | n.8G > C^ homozygous | | | |  |
| F362.1 | 2 Y | | n.20C > T/n.*5C > G | | | |  |
| F362.2 | 2 Y | | n.20C > T/n.*5C > G | | | |  |
| F362.3 | 2 Y | | n.20C > T/n.*5C > G | | | |  |
| F414 | 25 Y | | n.61A > G/n.*1C > T | | | |  |
| F426.1 | <6 Mo | | n.81G > A/n.*5C > G | | | |  |
| F426.2 | <6 Mo | | n.81G > A/n.*5C > G | | | |  |
| F433 | 54 Y | | n.20C > T/n.5*C > G | | | |  |
| F445 | 6 Mo | | n.127C > G/n.*5C > G | | | | Yes |
| F446 | 10 Mo | | n.*5C > G homozygous | | | |  |
| F454.1 | 14 Y | | *n.-54_-49del*/n.*5C > G | | | |  |
| F454.2 | 11 Y | | *n.-54_-49del*/n.*5C > G | | | |  |
| F465 | 15 Mo | | n.60_61insT/n.82A > G | | | |  |
| F521.1 | <12 Mo | | n.104G > A/n.131C > G | | | |  |
| F521.2 | 1 Y | | n.104G > A/n.131C > G | | | |  |
| F551 | <1 Y | | n.127C > G/n.*9C > T | | | | Yes |
| F564 | 9 Mo | | n.126C > T/n.*1C > T | | | | Yes |
| F641 | 4 Mo | | n.61A > G/n.*10G > T | | | |  |
| F691 | 10 Mo | | n.58A > G/n.*9C > T | | | |  |
| F730 | 2 Mo | | n.81G > A/n.*10G > T | | | |  |
| F766 | 12 Y | | n.3C > A/n.42G > A^b^ | | | |  |
| F780.1 | 3 Mo | | n.131C > G/n.*1C > T | | | |  |
| F780.2 | 6 Y | | n.131C > G/n.*1C > T | | | |  |
| F819.1 | 12 Y | | *n.-70_n.?76del*/n.*1C > T^^^^ | | | |  |
| F819.2 | 5 Y | | *n.-70_n.?76del*/n.*1C > T^^^^ | | | |  |
| F856 | 11 Y | | n.24C > T^b^/n.*10G > T | | | |  |
| F906 | 1 Y | | n.39G > C^a^/n.103G > A | | | | Yes |
| F1127 | <6 Mo | | n.100T > G/n.*9C > T | | | | Yes |
| F1288 | 3 Y | | n.59T > G/n.*9C > T | | | |  |
| F1424 | 6 Mo | | *n.-6G > A^a^*/n.130T > C | | | | Yes |
| F1445 | 8 Y | | n.3C > T/n.81G > A | | | |  |
| F1722 | 42 Y | | n.*1C > T/n.*5C > G | | | | Yes |
| F1954 | 4 Mo | | n.39G > T^a^ homozygous | | | |  |
| F1968 | 22 Y | | *n.-6G > A^a^*/n.131C > G | | | |  |
| F2036** | 54 Y | | n.81G > A/*n.*10G > C*^a^ | | | |  |
| F2054 | <6 Mo | | n.20C > T/n.*10G > T | | | |  |
| F2089 | 21 Y | | n.81G > A/n.*9C > T^^^ | | | |  |
| F2143 | 59 Y | | n.64G > A/n.*9C > T | | | | Yes |
| F2243 | 67 Y | | *n.-6G > A^a^*/n.74G > A | | | |  |
| F2380 | 2 Mo | | n.57G > T/n.*5C > G | | | |  |
| F2427 | 25 Y | | n.72A > G/n.*9C > T | | | |  |
| F2474 | 42 Y | | n.73T > G/n.*5C > G | | | |  |
| F2480 | 12 Y | | n.72A > G/n.*10G > T | | | |  |
| F2494 | 12 Y | | n.3C > T/n.57G > A | | | |  |
| F2689 | 2 Y | | n.39G > C^a^/n.75A > C | | | |  |
| F2737 | 3 W | | n.56dupT/n.*5C > G | | | |  |
| F2816 | 48 Y | | n.58A > G/n.*9C > T | | | |  |
| F2871 | 9 Y | | *n.-54_-49del*/n.*9C > T | | | |  |
| N8058 | 28 Y | | *n.-6G > A^a^*/n.131C > A | | | |  |
| N11301 | 15 Y | | n.3C > A/n.20C > T | | | |  |
| DIF_1 | 37 Y | | n.60G > C/n.*9C > T | | | |  |
| Hild_1 | 1 Y | | n.8G > C homozygous | | | |  |
| Iwama et al. 2017 | 45 Y | | | n.39G > C^a^/n.72A > G | |  | |
|  | 53 Y | | | n.39G > C^a^/n.118T > G | | Yes | |
|  | 11 Mo | | | n.39G > C^a^/n.103G > A | | Yes | |
|  | n.d. | | | n.3C > T/n.19C > G | |  | |
|  | 1 Mo | | | n.39G > C^a^/n.72A > G | |  | |
|  | 9 Y | | | n.3C > T/n.24C > T^b^ | |  | |
|  | 37 Y | | | n.39G > C^a^/n.61A > T | | Yes | |
|  | 1 Y | | | n.39G > C^a^ homozygous | |  | |
| Iwasaki et al. 2017 | 4 Y | | | n.39G > C^a^/n.117C > G | | Yes | |
| Jin et al. 2018 | n.d. | | | n.82A > G/n.*10G > A | |  | |
| Taglia et al. 2018 | 2 Mo | | | n.57G > A/n.*10G > T | |  | |
| Pessoa 2018 | 24 Y | | | n.61A > G/n.*1C > T | |  | |
| Hermens et al. 2018 | 11 Mo | | | n.24C > A^a^/n.82A > G | |  | |
| Osman et al. 2020 | 30 Y | | | n.3C > T/n.72A > G | |  | |
|  | 28 Y | | | n.74G > T/n.*9C > T | |  | |
|  | 11 Y | | | n.72A > G/n.92C > T | |  | |
|  | 18 Y | | | n.39_40insT^a^/n.72A > G | |  | |
|  | 60 Y | | | n.131C > T/n.*9C > T | |  | |

**Table S2 A summary of LCC genotypes**^1–7^**.** Hypomorphic and null mutations are coloured black and red respectively, and alleles of unknown function in grey. Italicised alleles were not assessed in the zebrafish assay. Whole gene deletions are presumed null, and the n.-54_-49del U8 variant has previously been reported to almost completely inactivate U8 promoter activity, indicating that it acts as a null mutation^8^.

^^^F344 is homozygous for both n.8G > C and n.113C > T.

^^^^F819: both children inherited a paternal deletion, but different maternally inherited mutations. The asymptomatic mother is compound heterozygous for n.*1C > T and n.*9C > T.

^^^^^F2089 also carried a n.119G > T in combination with n.*9C > T.

*F330 originally reported as n.8G > A/n.82A > G. Sanger sequencing analysis determined n.8G > A co-segregates with null n.82A > G on the same allele. On the other allele n.25C > T is the only other candidate rare variant.

**F2036 originally reported as n.81G > A/n.*10G > T. Sanger sequencing analysis determined the correct genotype is n81G > A/n.*10G > C.

^a^denotes U8 mutants that are detected in homozygous form in patients with LCC (n.39G > C, n.39G >T), or the gnomAD v4.1.0 database (n.-6G > A, n.39_40insT), or co-occur with known null alleles in patients with LCC (n.24C > A, n.25C >T, n.*10G > C), indicating hypomorphic function.

^b^denotes U8 mutants whose functionality is uncertain due to evidence that the affected nucleotide localises to regions of human U8 that are not conserved at the sequence level in zebrafish.

| **Crow cohort** | | | |
| --- | --- | --- | --- |
| **Patient** | **Genotype** | **n.105 allelic status** | |
| F172 | n.56dupT/n.*10G > T | | Heterozygous. n.105T segregates with n.*10G>T |
| F278 | n.8G > C/n.75A > G | | Homozygous T |
| F281 | n.3C > T/n.81G > C | | Homozygous T |
| F285 | n.57G > A/n.*5C > G | | Homozygous T |
| F309 | n.3C > T/n.131C > G | | Heterozygous. n.105T segregates with n.3C>T |
| F330 | *n.25C > T*^a^/n.82A > G | | Heterozygous. n.105T segregates with n.25C>T |
| F331.1 | n.72A > G/n.*10G > T | | Heterozygous. n.105T segregates with n.*10G>T |
| F331.2 | n.72A > G/n.*10G > T | | Heterozygous. n.105T segregates with n.*10G>T |
| F334 | n.82A > G/n.*10G > T | | Homozygous T |
| F337 | n.2T > C/n.58dup | | Homozygous T |
| F343 | *n.-7_22dup*/n.*5C > G | | Homozygous T |
| F344 | n.8G > C^^^ homozygous | | Homozygous A |
| F362.1 | n.20C > T/n.*5C > G | | Homozygous T |
| F362.2 | n.20C > T/n.*5C > G | | Homozygous T |
| F362.3 | n.20C > T/n.*5C > G | | Homozygous T |
| F414 | n.61A > G/n.*1C > T | | Homozygous T |
| F426.1 | n.81G > A/n.*5C > G | | Homozygous T |
| F426.2 | n.81G > A/n.*5C > G | | Homozygous T |
| F433 | n.20C > T/n.5*C > G | | Homozygous T |
| F445 | n.127C > G/n.*5C > G | | Homozygous T |
| F446 | n.*5C > G homozygous | | Homozygous T |
| F454.1 | *n.-54_-49del*/n.*5C > G | | Heterozygous. n.105T segregates with n.*5C>G |
| F454.2 | *n.-54_-49del*/n.*5C > G | | Heterozygous. n.105T segregates with n.*5C>G |
| F465 | n.60_61insT/n.82A > G | | Homozygous T |
| F521.1 | n.104G > A/n.131C > G | | Heterozygous. n.105T segregates with n.104G>A |
| F521.2 | n.104G > A/n.131C > G | | Heterozygous. n.105T segregates with n.104G>A |
| F551 | n.127C > G/n.*9C > T | | Homozygous T |
| F564 | n.126C > T/n.*1C > T | | Homozygous T |
| F641 | n.61A > G/n.*10G > T | | Not assessed |
| F691 | n.58A > G/n.*9C > T | | Homozygous T |
| F730 | n.81G > A/n.*10G > T | | Heterozygous. n.105T segregates with n.*10G>T |
| F766 | n.3C > A/n.42G > A^b^ | | Homozygous T |
| F780.1 | n.131C > G/n.*1C > T | | Homozygous T |
| F780.2 | n.131C > G/n.*1C > T | | Homozygous T |
| F819.1 | *n.-70_n.?76del*/n.*1C > T^^^^ | | Homozygous T |
| F819.2 | *n.-70_n.?76del*/n.*9C > T^^^^ | | Homozygous T |
| F856 | n.24C > T^b^/n.*10G > T | | Homozygous T |
| F906 | n.39G > C^a^/n.103G > A | | Heterozygous. n.105T segregates with n.103G>A |
| F1127 | n.100T > G/n.*9C > T | | Homozygous T |
| F1288 | n.59T > G/n.*9C > T | | Not assessed |
| F1424 | *n.-6G > A^a^*/n.130T > C | | Homozygous T |
| F1445 | n.3C > T/n.81G > A | | Homozygous T |
| F1722 | n.*1C > T/n.*5C > G | | Homozygous T |
| F1954 | n.39G > T^a^ homozygous | | Not assessed |
| F1968 | *n.-6G > A^a^*/n.131C > G | | Heterozygous. n.105T segregates with n.-6G>A |
| F2036 | n.81G > A/n.*10G > C | | Homozygous T |
| F2054 | n.20C > T/n.*10G > T | | Homozygous T |
| F2089 | n.81G > A/n.*9C > T^^^ | | Heterozygous. n.105A segregates with n.*9C>T |
| F2143 | n.64G > A/n.*9C > T | | Homozygous T |
| F2243 | *n.-6G > A^a^*/n.74G > A | | Homozygous T |
| F2380 | n.57G > T/n.*5C > G | | Heterozygous. n.105T segregates with n.*5C>G |
| F2427 | n.72A > G/n.*9C > T | | Homozygous T |
| F2474 | n.73T > G/n.*5C > G | | Homozygous T |
| F2480 | n.72A > G/n.*10G > T | | Heterozygous. n.105T segregates with n.*10G>T |
| F2494 | n.3C > T/n.57G > A | | Homozygous T |
| F2689 | n.39G > C^a^/n.75A > C | | Not assessed |
| F2737 | n.56dup/n.*5C > G | | Not assessed |
| F2816 | n.58A > G/n.*9C > T | | Heterozygous. n.105T segregates with n.*9>T |
| F2871 | *n.-54_-49del*/n.*9C > T | | Not assessed |
| N8058 | *n.-6G > A^a^*/n.131C > A | | Not assessed |
| N11301 | n.3C > A/n.20C > T | | Not assessed |
| DIF_1 | n.60G > C/n.*9C > T | | Not assessed |
| Hild_1 | n.8G > C homozygous | | Not assessed |

**Table S3** **Summary of allelic variation at position n.105 of U8 in patients from a previously reported cohort of patients with LCC**^1^ Individuals in whom n.105A segregates with a known hypomorphic mutation are highlighted in green. All other patients are unable to methylate n.105 on hypomorphic alleles of U8. Hypomorphic and null mutations are coloured black and red respectively, and alleles of unknown function in grey. Italicised alleles were not assessed in the zebrafish assay. Whole gene deletions are presumed null, and the n.-54_-49del U8 variant has previously been reported to almost completely inactivate U8 promoter activity^2^, indicating that it acts as a null mutation

^^^F344 is homozygous for both n.8G > C and n.113C > T.

^^^^F819: both children inherited a paternal deletion, but different maternally inherited mutations. The asymptomatic mother is compound heterozygous for n.*1C > T and n.*9C > T.

^^^^^F2089 also carries a n.119G > T that co-segregates with n.*9C > T.

^a^denotes U8 mutants that are detected in homozygous form in patients with LCC (n.39G > C, n.39G >T), or the gnomAD v4.1.0 database (n.-6G > A, n.39_40insT), or co-occur with known null alleles in patients with LCC (n.24C > A, n.25C > T, n.*10G > C), indicating hypomorphic function.

^b^denotes U8 mutants whose functionality is uncertain due to evidence that the affected nucleotide localises to regions of human U8 that are not conserved at the sequence level in zebrafish.

1.	Crow, Y. J. *et al.* Leukoencephalopathy with calcifications and cysts: Genetic and phenotypic spectrum. *Am J Med Genet A* (2020) doi:10.1002/ajmg.a.61907.

2.	Jenkinson, E. M. *et al.* Mutations in SNORD118 cause the cerebral microangiopathy leukoencephalopathy with calcifications and cysts. *Nat Genet* (2016) doi:10.1038/ng.3661.

| **Comparison** | **p-value** | **p-value**  **adjusted** | **Significant** |
| --- | --- | --- | --- |
| Intramolecular duplex vs  Protein interaction domains | 0.001148 | 0.005739 | Yes, increased allelic variation in the intramolecular duplex |
| Stem-loop 1 vs Protein interaction domains | 0.00308 | 0.003076 | Yes, increased allelic variation in stem-loop 1 |
| Stem-loop 3 vs Protein interaction domains | 0.013090 | 0.032724 | Yes, increased allelic variation in stem-loop 3 |
| Loop 3 vs Protein interaction domains | 0.002078 | 0.006925 | Yes, increased allelic variation in loop 3 |
| Loop 3 vs intramolecular duplex | 0.429438 | 0.429438 | No, allelic variation is not significantly different in these regions of U8 |
| Loop 3 vs Stem-loop 1 | 0.406313 | 0.429438 | No, allelic variation is not significantly different in these regions of U8 |

**Table S4 Mann-Whitney U-test (FDR corrected) of allelic variation within the U8 snoRNA**

Protein interaction domains of U8 encompass nucleotides 55-61, 67-90, 130-134. Intramolecular duplex specific to precursor U8 encompasses 1-15, 137-161. Stem-loop 1 of U8 encompasses nucleotides 29-43. Stem-loop 3 of U8 encompasses nucleotides 96-124. Loop 3 of U8 encompasses nucleotides 105-115.
